# A vasopressin circuit that modulates sex-specific social interest and anxiety-like behavior in mice

**DOI:** 10.1101/2023.11.06.564847

**Authors:** Nicole Rigney, Elba Campos-Lira, Matthew K. Kirchner, Wei Wei, Selma Belkasim, Rachael Beaumont, Sumeet Singh, Geert J. de Vries, Aras Petrulis

## Abstract

One of the largest sex differences in brain neurochemistry is the male-biased expression of the neuropeptide arginine vasopressin (AVP) within the vertebrate social brain. Despite the long-standing implication of AVP in social and anxiety-like behavior, the precise circuitry and anatomical substrate underlying its control are still poorly understood. By employing optogenetic manipulation of AVP cells within the bed nucleus of the stria terminalis (BNST), we have unveiled a central role for these cells in promoting social investigation, with a more pronounced role in males relative to females. These cells facilitate male social investigation and anxiety-like behavior through their projections to the lateral septum (LS), an area with the highest density of sexually-dimorphic AVP fibers. Blocking the vasopressin 1a receptor (V1aR) in the LS eliminated stimulation-mediated increases in these behaviors. Together, these findings establish a distinct BNST AVP → LS V1aR circuit that modulates sex-specific social interest and anxiety-like behavior.

**Significance Statement:** The function of sex differences in the brain is poorly understood. Here we test the function of one of the most consistently found sex differences in vertebrate brains, the male-biased vasopressin projections from the bed nucleus of the stria terminalis. Using optogenetic techniques, we demonstrate that these cells and their projection to the lateral septum are much more important in driving male than female social investigation. These studies make a strong contribution to understanding how sexually dimorphic circuitry controls social behavior.

## Introduction

Dysfunction in social behavior and communication prominently features in psychopathologies such as autism, schizophrenia, and social anxiety <u>(1, 2)</u>. These disorders show marked sex differences, with, for example, autism and schizophrenia being more prevalent in males and social anxiety more prevalent in females <u>(2, 3)</u>. Understanding the nature of these differences may facilitate developing new avenues for treating these disorders. A reasonable hypothesis is that differences in neural circuitry that control social behavior contribute to sex differences in dysfunctions of social behavior and communication. A problem is that there is no clear understanding of how sex differences in the neurobiology of social behavior occur, whether normal or disordered <u>(4)</u>.

A promising system to study in this regard is the vasopressin innervation of the brain. AVP has been repeatedly implicated in modulation of social behaviors in sex-different ways <u>(5–9)</u> and is an important modulator for both animal <u>(10)</u> <u>(6)</u> and human sociality <u>(11)</u>. In humans, AVP has been implicated in psychopathology <u>(12)</u>. For example, both variations in the vasopressin V1a receptor (V1aR) gene and AVP serum levels are associated with autism <u>(13)</u>. Pharmacological studies indicate that AVP acts on various brain regions that regulate social communication <u>(14)</u>, aggression <u>(15)</u> maternal care <u>(16)</u>, pair bonding <u>(17)</u>, cognition <u>(18)</u>, and social recognition <u>(5)</u>. Additionally, AVP contributes to anxiety-related behavior <u>(6)</u>, passive avoidance <u>(19)</u> as well as in the evaluation of stressful situations <u>(20)</u>.

The anatomy of AVP projections suggests that AVP control of social behavior is complex. Central AVP projections stem from a diverse set of cell groups, the most prominent of which are found in the paraventricular of the hypothalamus, the suprachiasmatic nuclei of the hypothalamus, and the bed nucleus of the stria terminalis (BNST) and medial amygdaloid nucleus (MeA), each of which project to distinct but sometimes overlapping brain areas <u>(21)</u>. The most sexually dimorphic AVP projections come from the BNST and MeA <u>(22)</u>. Male rodents, for example, have about twice as many AVP cells as females in the BNST and their projections to areas such as the lateral septum (LS) are denser as well <u>(21, 23)</u>. This is one largest and most evolutionarily conserved sex-different systems in the vertebrate brain <u>(14, 22, 24)</u>.

Most studies that implicated specific AVP projections in the control of social behavior did so indirectly, e.g., by pharmacological blockade of AVP receptors in specific areas in the brain <u>(6, 25)</u>. As it is nearly impossible to restrict the effects of such manipulations to AVP projections from just one source, it remained unclear specifically which of the above cell groups contribute to AVP effects of social behavior. Recently, we have started to systematically test the role of specific AVP cell groups in social behavior by blocking AVP expression in specific cell groups <u>(26)</u> or by lesioning AVP cells using viral vector approaches <u>(27–29)</u>. Both of these manipulations blocked social behavior in male but not in female mice. As both of them chronically impaired BNST AVP function, we cannot rule out that BNST projections acutely control social behavior in females. Moreover, we do not know the behavioral effects of activating this cell population. Neither do we know which BNST AVP projections modulate social behavior.

In this study we address all of these questions by showing that exposure to social stimuli acutely activates fos expression in BNST AVP cells. We also show that optogenetic inhibition of those cells activity reduced same sex social investigation in males but not in females, whereas optogenetic stimulation of those cells increased same and opposite sex social investigation, but more so in males than in females. Finally, we show that activating BNST AVP projections to the LS, one of the most conspicuous targets of BNST AVP cells <u>(21, 30)</u> as well as a site where AVP has consistently be shown to influence social behavior <u>(6, 9, 31)</u> had an overall inhibitory effect on septal neuronal activity while increasing social investigation in males, effects that could be blocked by an AVP V1a receptor antagonist.

## Results

### BNST AVP cells are activated during social investigation

AVP cells in the BNST are activated during copulatory and aggressive interactions in male mice <u>(32)</u>, but it is unclear whether these cells are also activated during social investigation. To answer this, we measured Fos expression within BNST AVP cells of males and females exposed to a caged male conspecific, female conspecific, or empty cages in a three-chamber apparatus. Male mice exposed to males or females had increased BNST AVP-Fos colocalization compared to males in an empty three-chamber apparatus (Figure 1a-c). Female mice showed similar Fos activation in BNST AVP cells (Figure 1a-c). However, females had less AVP-Fos colocalization in the BNST in both social conditions compared to males, and females had less overall BNST Fos. These results suggest that BNST AVP cells encode social cues and may drive social approach and communicative behaviors in both sexes, but less so in females.

**Figure 1.**
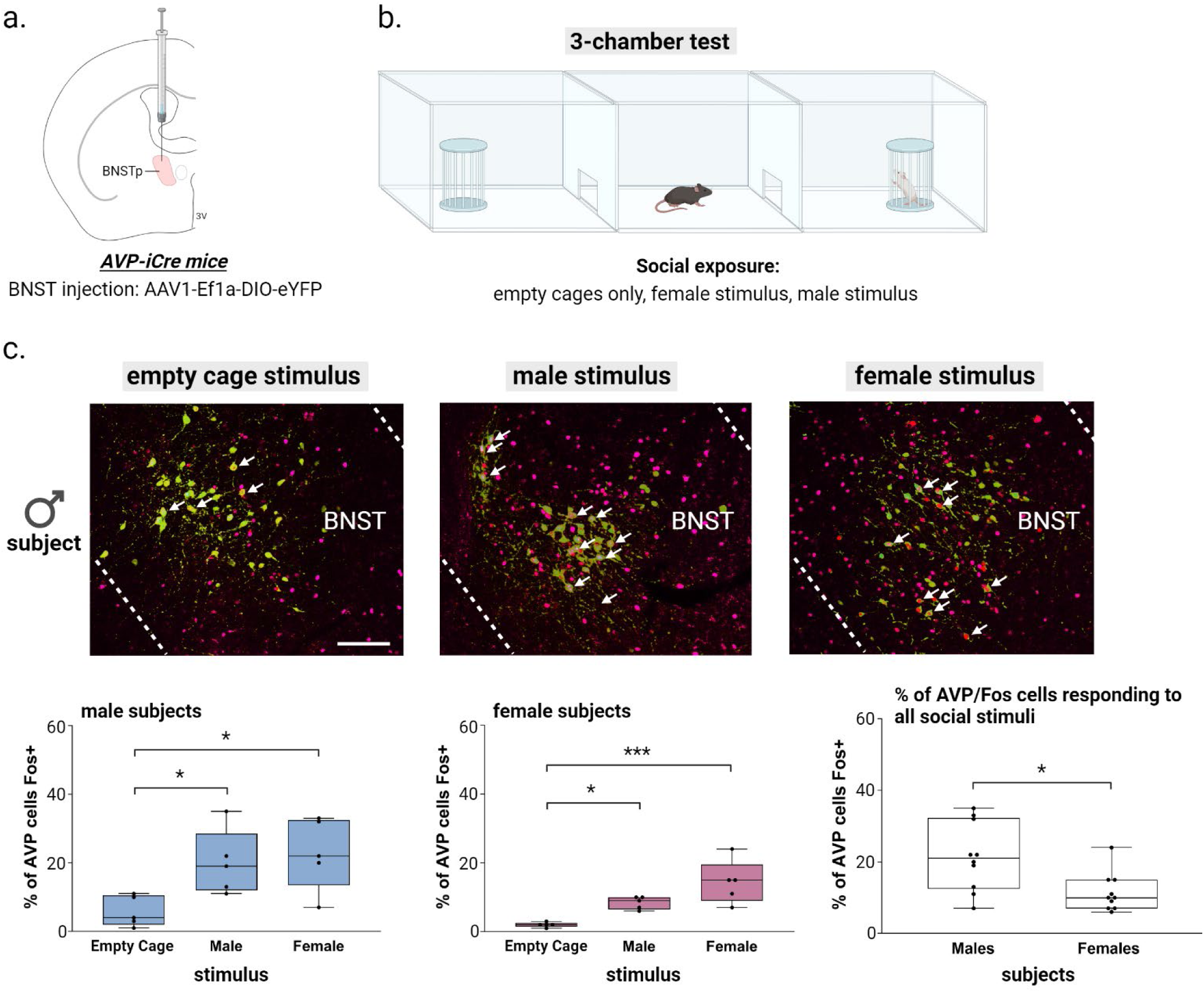
BNST AVP cell and Fos colocalization during social exposure in the 3-chamber test (a) BNST injection site and virus. (b) Three-chamber social testing. (c) (top) Example images of merged BNST-AVP cells (green) and Fos+ cells (magenta). (bottom) Boxplots of the percentage of AVP cells colocalized with Fos in each 3-chamber stimulus condition (empty cage, male, female) for male and female subjects. A Two-way ANOVA revealed a significant effect when comparing activated BNST AVP cells with the type of stimulus received (F(1,24) = 12.89, p = 0.00016, η2 = 0.8), a significant effect when comparing the sex of the subjects (F(1,24) = 10.69, p = 0.003, η2 = 0.62), yet there was no interaction between sex and stimulus type. A post hoc ANOVA followed by Bonferroni-corrected pairwise tests showed that male mice exposed to male or female conspecifics had increased BNST AVP-Fos colocalization compared to males kept in an empty three-chamber apparatus containing empty stimulus cages (F(1,12) = 5.8, p = 0.017, η2 = 0.4; male vs. empty cage: p = 0.05, female vs. empty cage: p = 0.025). Similarly, female mice also showed Fos activation to social cues in BNST AVP cells (F(1,12) = 13.23, p = 0.0009, η2 = 0.69; male vs. empty cage: p = 0.05, female vs. empty cage: p = 0.0007). Mean ± SEM data represented. Dots indicate individual data points. White arrows indicate Fos and AVP cell colocalization. Scale bar = 25 µm. *p < 0.05, ** p < 0.01, ***p < 0.001.

### Inhibiting BNST AVP cells reduces male’s social investigation of other males, without affecting social communicative behaviors

Deletion of BNST AVP cells alter male, but not female, social investigation and communication <u>(27)</u>. As compensatory changes may have taken place after deletion, we cannot exclude the possibility that these cells affect social behavior in both sexes. To address this question, we tested whether acute inhibition of BNST AVP cells using an optogenetic approach (Figure 2a-b) influenced male and female social investigation and male-biased social communication behaviors such as urine marking <u>(33)</u> and ultrasonic vocalizations <u>(34)</u>. To confirm that light application inhibits AVP BNST cells expressing the inhibitory blue-light opsin stGTACR2, we injected adeno-associated viruses (AAV) expressing Cre-dependent stGTACR2-FusionRed into the BNST of adult male and female AVP-iCre+ mice and, after 3 weeks, performed patch-clamp recordings of BNST FusionRed-tagged AVP cells under blue-light stimulation. We found that continuous (10 and 90 second) blue light reliably inhibited current-injected BNST cells in both males and females (Figure 2c).

**Figure 2.**
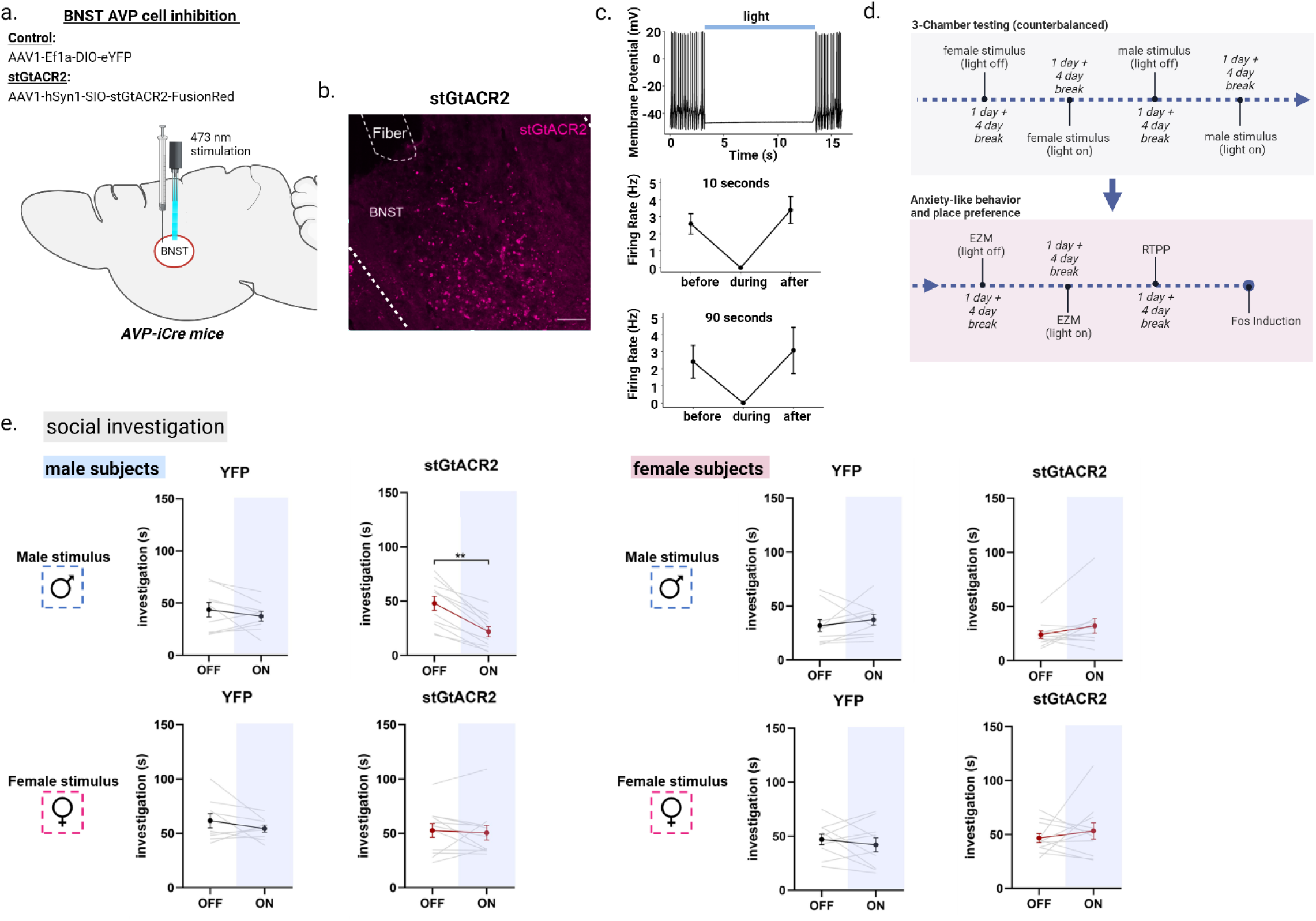
Optogenetic inhibition of AVP-BNST cells decreases male-male social investigation. (a) Bilateral BNST injection and fiber implantation site; coordinates: DV: -4.4, AP: +0.15, ML: ±0.8; modified from Paxinos and Franklin (2012). (b) Example image of BNST AVP cells infected with inhibitory stGtACR adeno-associated virus (FusionRed) (c) representative trace from whole-cell current-clamp recording of stGTACR2-expressing BNST cell silenced by light application at 10Hz for 10 seconds and firing rate of stGTACR2-expressing BNST cell suppressed by the blue light for 10 seconds (n=8 cells) and 90 seconds (n=9 cells). (d) Experimental timeline. All subjects were tested within the 3-chamber apparatus on 4 separate days (4-day break in between) with all stimuli and light conditions counterbalanced. The same subjects were further tested within the elevated-zero maze (EZM) (counterbalanced) followed by a real-time place preference test (RTPP) and Fos induction. (e) Social investigation (in seconds) by male and female subjects during the three-chamber test (male subjects: YFP, n=9 and stGtACR2, n=11 female subjects (YFP, n=10 and stGtACR2, n=11) during light-OFF and light-ON conditions, counterbalanced. Blue light inhibition (ON) of AVP-BNST cells in stGtACR2 males significantly decreased time spent investigating male stimuli, but not female stimuli, compared to investigation during light-OFF condition (Mixed model ANOVA, treatment*light*stimulus interaction (F(1,37) = 4.11, p = 0.05, η^2^ = 0.6; *post hoc*: p = 0.001). Blue light inhibition (ON) of AVP-BNST cells in stGtACR2 females did not affect time spent investigating male or female stimuli. Light stimulation did not affect investigation times of YFP male and female subjects to either stimulus type (female or male). Bonferroni *post hoc* tests were used. Each point and horizontal line represent individual within-subject data. Overlapping data are represented as one point/line. Scale bar = 50 µm. **p<0.01.

After confirming *ex vivo* efficacy of stGTACR2-mediated inhibition in BNST AVP cells, we bilaterally injected AAVs that Cre-dependently expressed stGTACR2-FusionRed or eYFP as a control in BNST AVP cells and bilaterally implanted optic fibers into the BNST of adult male and female AVP-iCre+ mice. Histological analysis indicated that FusionRed expression was limited to the BNST (Figure 2b), and males had approximately 40-50% more cells labeled than females, reflecting the sex difference in the number of AVP-expressing cells in these mice that we found earlier <u>(27)</u>. After 3 weeks, subjects were introduced into a three-chamber apparatus containing either a caged male conspecific or caged female conspecific (each paired with an empty cage) and were tested twice in each condition: once with delivery of constant blue light (473 nm, 5-6 mW light power) and once with no light delivery (Figure 2d). Upon light delivery, stGTACR2 males significantly decreased their time spent investigating male stimuli, compared to investigation levels during the light-off condition; investigation of female conspecifics was unaltered (Figure 2d). In all conditions, investigation of the empty cage was unaltered by light stimulation, indicating no general changes in investigation or activity (Figure S1). In contrast, levels of social investigation by stGTACR2 females did not differ between light-on and light-off conditions (Figure 2e). Social investigation by control eYFP males or females also did not differ during light-on and light-off trials. Moreover, inhibition of BNST AVP cells did not affect social communicative behaviors (i.e., urine marking and ultrasonic vocalizations) in males (Figure S2) or in females, which produced little to no urine marks and vocalizations in any condition (data not shown).

As changes in social investigation may be explained by changes in anxiety or valence processing, both of which are affected by central AVP <u>(35, 36)</u>, we tested whether inhibiting BNST AVP cells altered these states. First, we observed that optogenetic inhibition of BNST AVP cells did not alter time spent in the open, anxiogenic, arms of the elevated zero maze (EZM) (Figure S3), indicating no obvious change in anxiety-like states. Second, we examined whether inhibiting BNST AVP cells is intrinsically rewarding or aversive using a real-time place preference test (RTTP). During this test stGTACR2 females, but not males, spent more time in the light-paired chamber compared to eYFP control females, suggesting that inhibition of BNST AVP cells was moderately rewarding to females (Figure S3).

### Exciting BNST AVP cells increases social investigation in both sexes as well as male social communication

As BNST AVP cells show increases in response to social cues (see above), we tested whether increasing BNST AVP cell activity would boost social investigation and communication. We first confirmed *ex vivo* that blue light stimulated AVP BNST cells of male and female AVP-iCre+ mice that Cre-dependently expressed the excitatory opsin ChR2-eYFP (Figure 3d). Increasing frequency of light stimulation increased activity in BNST AVP cells from both males and females (Figure 3d). However, stimulation beyond 10 Hz led to decreased spike fidelity (Figure 3d). Consequently, we used 10 Hz stimulation, a frequency known to release neuropeptide-containing vesicles <u>(37)</u>, for all subsequent *in vivo* procedures.

**Figure 3.**
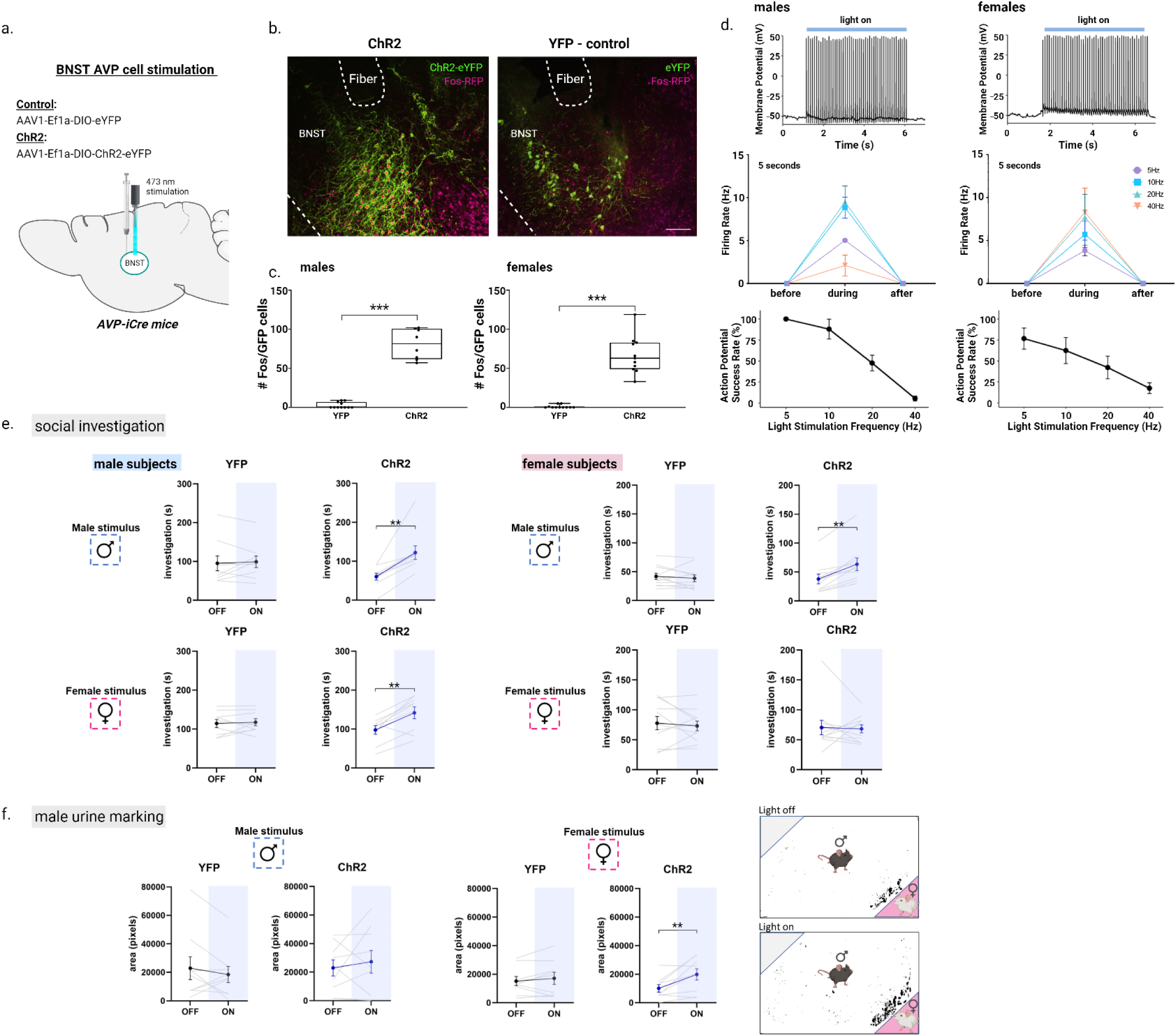
Optogenetic activation of AVP-BNST cells increases social investigation in both sexes and male urine marking toward female stimuli. (a-b) Bilateral BNST injection and fiber implantation site; coordinates: DV: -4.4, AP: +0.15, ML: ±0.8; modified from Paxinos and Franklin (2012). (b) Example images of merged BNST-AVP cells infected with either the excitatory ChR2 adeno-associated virus (ChR2) or YFP control virus, both colocalized with Fos+ cells (magenta). (c) Boxplots of the number of BNST AVP ChR2/YFP cells colocalized with Fos. Blue light stimulation robustly increased the number of BNST AVP labeled cells colocalized with Fos (F (1,35) = 234.17, p < 0.000001; η^2^ = 0.87). (d) Representative trace from whole-cell current-clamp recording of ChR2-mCherry-expressing cells activated by light application at 10Hz for 5 seconds and the response to 10ms pulse delivered at 5, 10, 20, 40Hz for 5 seconds, with the probability of action potential (AP) in response to an individual pulse (action potential success rate in male and female subjects (e) Social investigation (in seconds) by male and female subjects during the three-chamber test (male subjects: YFP, n=9 and ChR2, n=9; female subjects (YFP, n=10 and ChR2, n=11) during light-OFF and light-ON conditions, counterbalanced. Blue light stimulation (ON) of AVP-BNST cells in ChR2 males significantly increased time spent investigating male and female stimuli compared to investigation during light-OFF condition (Mixed model ANOVA, treatment*light*sex interaction, F (1,36) = 7.02, p = 0.012; η^2^ = 0.5; *post hoc*: p = 0.002 (male stimuli), p = 0.004 (female stimuli). Blue light stimulation (ON) of AVP-BNST cells in ChR2 females significantly increased time spent investigating male stimuli compared to investigation during light-OFF condition (*post hoc*: p = 0.002 (male stimuli). Light stimulation did not affect investigation times of YFP male and female subjects to either stimulus type (female or male). (f) Total area of urine marking by male subjects during the three-chamber test. Blue light stimulation (ON) of AVP-BNST cells in ChR2 males significantly increased urine marking in the presence of a female stimulus (Mixed model ANOVA, treatment*light*sex interaction, (F (1,36) = 4.5, p = 0.04; η^2^ = 0.4; *post hoc*: p = 0.009). Bonferroni *post hoc* tests were used. Each point and horizontal line represent individual within-subject data. Overlapping data are represented as one point/line. **p<0.01.

We then bilaterally injected AAVs into the BNST of male and female AVP-iCre+ mice that Cre-dependently expressed ChR2-eYFP or eYFP, as a control, in BNST AVP cells and bilaterally implanted optic fibers above the injection sites (Figure 3b). Histological analysis indicated that eYFP expression was limited to the BNST and males had about 50% more cells labeled than females, again reflecting the sex differences found in BNST AVP expression <u>(22, 27)</u>.. Subjects were then allowed to investigate caged male or caged female conspecifics (each paired with an empty cage) twice in a three-chamber apparatus: once during delivery of pulsed blue light (20 ms pulse width, 473 nm, 5-6 mW light power) and once with no light delivery. Blue-light activation of BNST AVP cells strongly increased the time that ChR2 males spent investigating male and female stimuli compared to their investigation during the light-off condition (Figure 3e). Surprisingly, activating BNST AVP cells in ChR2 females also increased their investigation time of male, but not female, stimuli (Figure 3e). We did not notice changes in locomotion or time spent investigating the clean stimulus cage (Figure S4). eYFP-expressing control males or females did not change their investigatory behavior across light-on and light-off trials (Figure 3e). Light stimulation of ChR2-expressing BNST AVP cells in males also increased male urine marking (total area) in the presence of female, but not male, stimuli (Figure 3f). This increase in marking was unrelated, at an individual level, to increases in social investigation (r^2^=0.27, p = 0.15), suggesting that marking was not directly linked to increased social interaction. In contrast to males, urine marking by females was low and was unaffected by optogenetic activation of BNST AVP cells. Control eYFP males and females did not change their marking behavior across light-on and light-off trials (Figure 3f). Activation of BNST AVP cells did not influence USV production (Figure S1). We also did not observe changes in anxiety-like behavior or real-time place preference across light-on/light-off conditions in either ChR2- or eYFP-expressing males and females, suggesting that the stimulation-induced changes in social investigation are not readily explained by changes in these behavioral states (Figure S5).

Following behavioral testing, a subset (n=34) of subjects were stimulated with blue-light, sacrificed, and processed for Fos/eYFP immunohistochemistry. We confirmed that blue-light stimulation of BNST AVP cells significantly increased Fos expression in these cells compared to eYFP controls (Figure 3b-c). Moreover, the number of excited BNST AVP cells (Fos/eYFP colocalized) in ChR2 males positively correlated with their time spent investigating female, but not male, stimuli (Figure S6). One explanation is that increased (artificial) recruitment of BNST AVP cells is needed for driving investigation of females, but that increased male-male investigation is, instead, triggered by a relatively low threshold of active cells.

### Activating AVP-BNST outputs to lateral septum increases social investigation and anxiety-like behavior in males, but not females

While more indirect approaches have supported the BNST as the major source of AVP innervation of the LS <u>(22)</u>, we recently confirmed that the BNST AVP cells send some of its strongest projections to the LS <u>(30)</u>. Here we tested whether these projections influence social behavior. First, to establish that exciting these projections indeed affects LS neuronal activity, we injected AAVs expressing Cre-dependent ChR2-mCherry into the BNST of adult male AVP-iCre+ mice and conducted patch-clamp recordings of LS cells adjacent to mCherry fibers under blue-light stimulation. We found that light stimulation of BNST AVP fibers within the LS led to a biphasic effect on LS cells, involving an initial excitation, followed by a more pronounced inhibition (Figure 4a-b). These results are broadly similar to those seen after exogenous application of AVP to LS *ex vivo* <u>(38)</u>. As optogenetic stimulation of BNST AVP → LS terminals likely releases neurotransmitters other than AVP, which may drive the complex pattern of response, we tested BNST AVP → LS stimulation in the presence or absence of a highly-selective V1aR antagonist. Intriguingly, we found that both excitatory and inhibitory responses in LS neurons induced by BNST AVP terminal stimulation were eliminated by the V1aR antagonist (Figure 4c). Overall, these results suggest that optogenetic stimulation of BNST AVP → LS terminals increases overall inhibition within the LS, an effect mediated by AVP acting on V1aRs.

**Figure 4.**
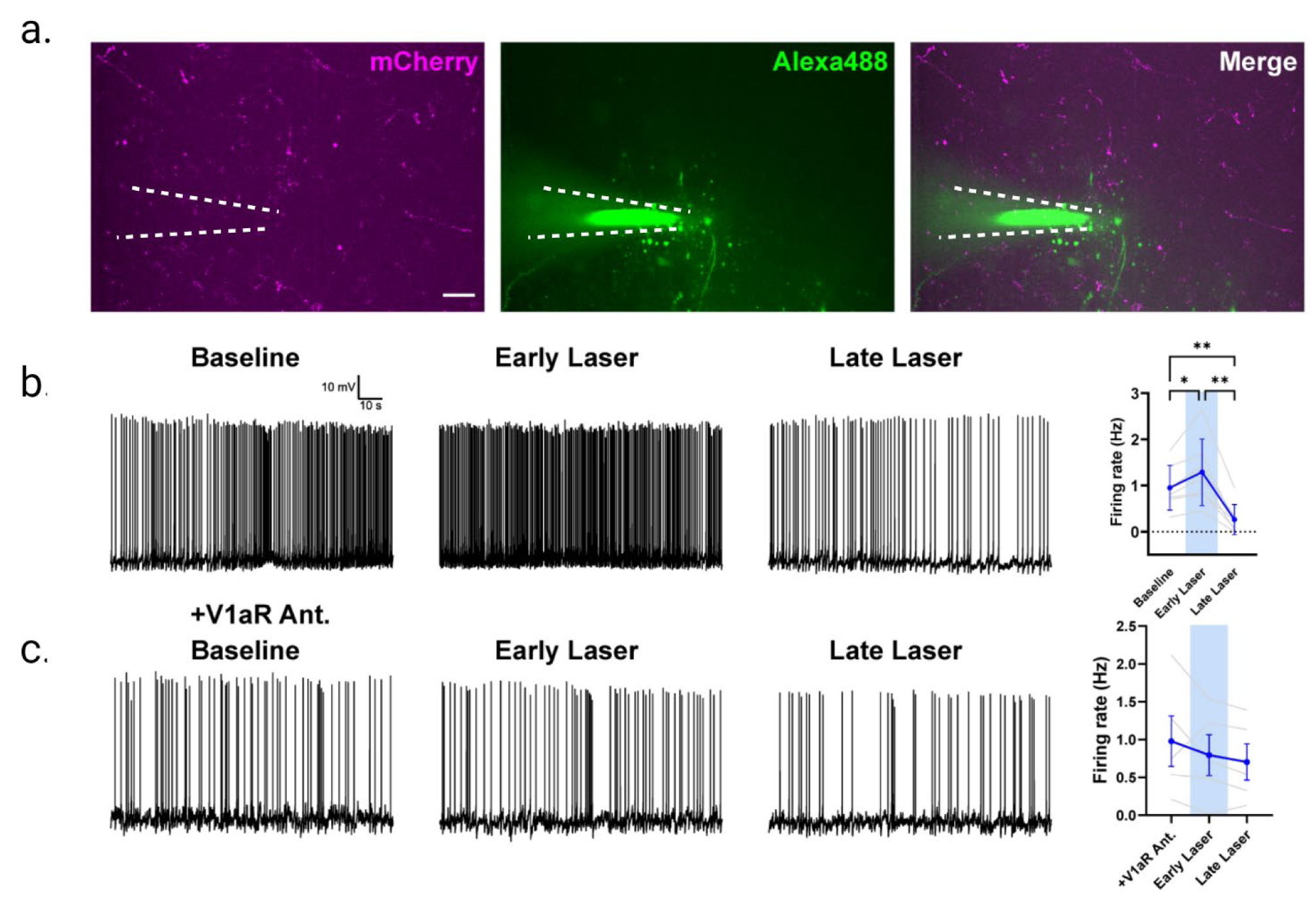
Optogenetic activation of AVP-BNST cell projections to the lateral septum (LS) modulates neurons in LS. (a) Visualization of mCherry reporter tagged to ChR2-AVP expressing fibers in the LS, originating from BNST (left) in addition to a patched neuron dialyzed with Alexa488 dye (middle). Merged image (right). (b) BNST AVP → LS terminal activation modulates LS neuron firing activity. (Baseline vs. Early Laser p <0.05, Baseline vs. Late Laser p <0.01, Early Laser vs. Late Laser p <0.01, one-way repeated measures ANOVA). (c) Bath application of V1aR antagonist (d(CH2)5[Tyr(Me)2,Dab5] AVP) blocks ChR2 modulation of LS neurons. (p >0.05, one-way repeated measures ANOVA). Scale bar= 25µm.

Next, we examined the behavioral effects of stimulating BNST AVP afferents to the LS in male and female mice. We bilaterally injected AAVs expressing Cre-dependent ChR2-eYFP or eYFP as a control into the BNST of adult male and female AVP-iCre+ mice and implanted fibers in the LS (Figure 5a). Blue light stimulation of BNST AVP → LS terminals in ChR2 males significantly increased their time spent investigating both male and female stimuli, compared to investigation during the light-off conditions; light stimulation did not affect their time spent investigating clean cages (Figure 5b; Figure S7). Blue light stimulation of BNST AVP → LS terminals did not affect social investigation in ChR2 females or eYFP subjects (Figure 5b). Similar to the effects of stimulating BNST AVP cells, blue light stimulation BNST AVP → LS terminals increased male urine marking. Unlike the prior experiment, BNST AVP → LS stimulation increased the number of marks, not the deposition area, in the presence of a female (Figure S8). However, as with BNST AVP stimulation, changes in scent marking levels were also not correlated with levels of social investigation (r^2^=0.01, p = 0.89), again suggesting that urine marking is not tightly linked to increased social investigation. As seen in prior experiments <u>(26, 27, 39)</u>, urine marking and USV production by females was low and unaffected by optogenetic stimulation (data not shown). In contrast to the lack of effect of BNST AVP cell stimulation (or inhibition) on anxiety-like behavior, stimulation of BNST AVP terminals in LS in ChR2 males significantly decreased time spent in the open arm of the EZM, indicating a possible anxiogenic effect (Figure 5c). However, increased anxiety-like behavior in subjects did not correlate with their increased social investigation (Figure S6). In females or eYFP controls, blue light stimulation had no effect on time spent in the open arm of the EZM (Figure 5c). Light stimulation was neither aversive nor rewarding in any group in the RTTP (Figure S5).

**Figure 5.**
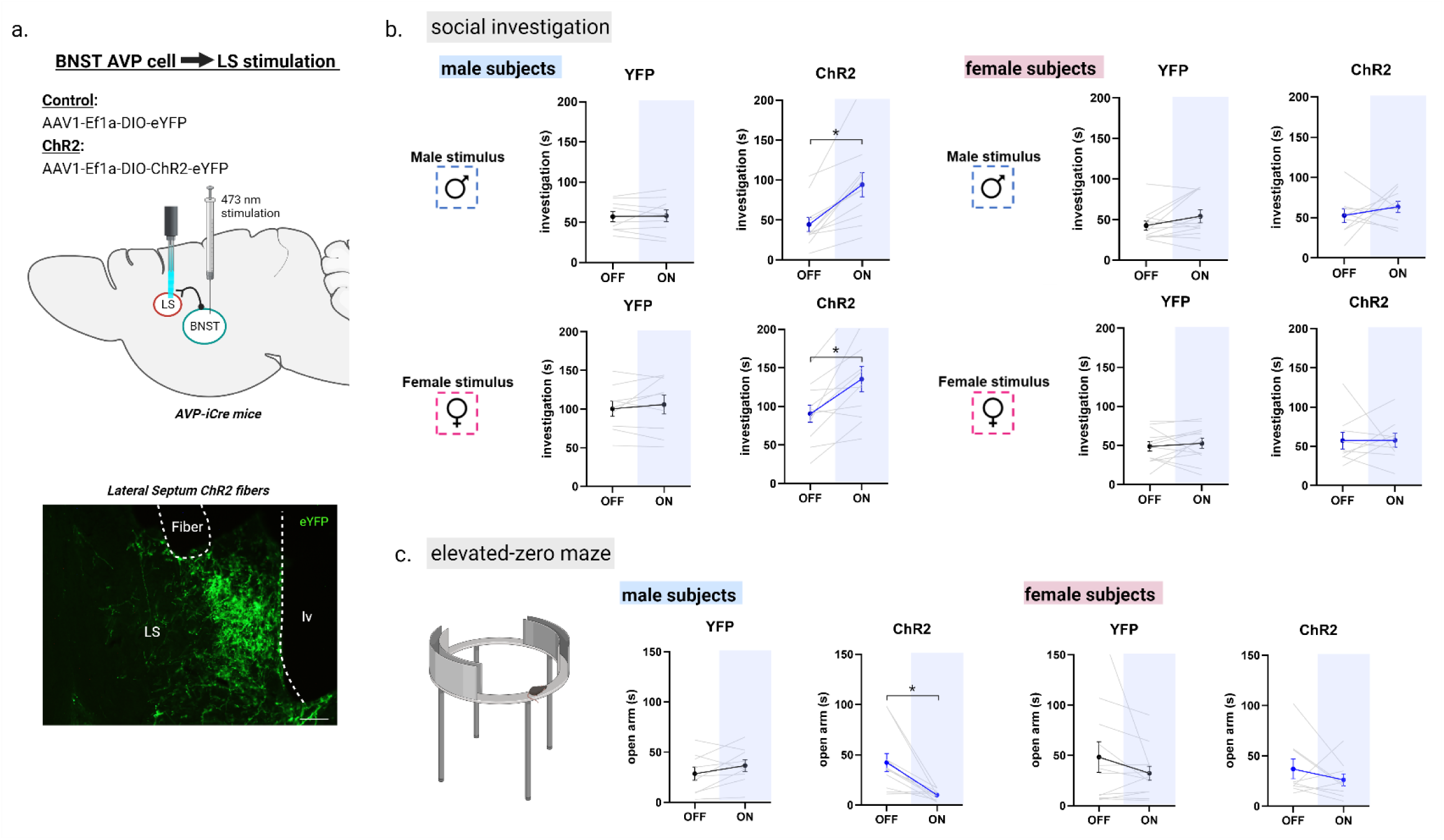
Optogenetic activation of AVP-BNST cell projections to the lateral septum (LS) increases social investigation and anxiety-like behavior in males, but not females. (a) Bilateral BNST injection of ChR2 adeno-associated virus (ChR2) and fiber implantation within the lateral septum (intermediate zone). (b) Social investigation (in seconds) by male and female subjects during the three-chamber test (male subjects: YFP, n=9 and ChR2, n=11; female subjects: YFP, n=12 and ChR2, n=9) during light-OFF and light-ON conditions, counterbalanced. Blue light stimulation (ON) of AVP-BNST-LS terminals in ChR2 males significantly increased time spent investigating male and female stimuli compared to investigation during light-OFF condition (Mixed model ANOVA, treatment*light interaction, (F (1,17) = 6.9, p = 0.01; η^2^ = 0.6; *post hoc*: p = 0.001 (male stimuli), p = 0.02 (female stimuli). Blue light stimulation (ON) of AVP-BNST LS terminals in ChR2 females did not affect investigation times compared to investigation during light-OFF condition. Light stimulation did not affect investigation times of YFP male and female subjects to either stimulus type (female or male). (c) Time spent in the open arm of the elevated-zero maze (EZM). Blue light stimulation (ON) of AVP-BNST-LS terminals in ChR2 males significantly decreased time spent in the open arm of the EZM. In females, blue light stimulation had no effect on time spent in the open arm of the EZM. (Mixed model ANOVA, treatment*light*sex interaction, (F (1,17) = 6.2, p = 0.024; η^2^ = 0.5; *post hoc*: p = 0.008). Bonferroni *post hoc* tests were used. Each point and horizontal line represent individual within-subject data. Overlapping data are represented as one point/line. *p<0.05.

### Increases in social investigation and anxiety-like behavior caused by BNST AVP → LS stimulation in males depend on the vasopressin 1a receptor

Although blocking V1aR action in LS reduces the impact of BNST AVP stimulation *ex vivo*, the behavioral effects of BNST AVP → LS stimulation may be due to the release of other neuroactive substances co-expressed in BNST AVP cells. Consequently, we tested if V1aR action was required for BNST AVP → LS optogenetic-mediated increases in male social investigation and anxiety-like behavior. Following Cre-dependent viral delivery of ChR2-eYFP to BNST AVP cells of male mice, cannulas with interchangeable fibers and fluid injectors were implanted in the LS to allow for dual delivery of V1aR antagonist (or saline control) and light stimulation in LS (Figure 6a). Two groups of ChR2 males were assigned to interact with either caged novel male or female conspecifics in the three-chamber apparatus. We tested subjects with their respective stimuli in four separate counterbalanced tests: light-off + saline infusion, light-on + saline infusion, light-off + V1aR antagonist infusion, light-on + V1aR antagonist infusion. Blue light stimulation of AVP BNST → LS terminals in both groups following saline injection significantly increased their time spent investigating male and female stimuli compared to their investigation during the light-off condition, replicating the behavioral effect of BNST AVP → LS stimulation (Figure 6b). In a test of non-social novel object investigation, a separate set of ChR2 male subjects (n=9) did not alter their investigation of non-social stimuli (toys) during BNST AVP → LS stimulation (Figure S9), indicating that optogenetic-mediated increases in male social investigation were limited to social targets rather than reflecting non-specific increases in investigatory behavior. Importantly, during pharmacological blockade of LS V1aR, BNST AVP → LS stimulation-mediated increases in social investigation of both male and female stimuli were eliminated (Figure 6b). We did note that V1aR antagonism by itself, during light-off conditions, reduced males’ investigation of both male and female stimuli compared to the saline control condition (Figure S10), further supporting the role of V1aR LS cells in social investigation. We were, however, unable to replicate the effect of BNST AVP → LS stimulation on male urine marking nor did we see any effect of our manipulation on USV production (Figure S8). LS V1aR antagonism by itself did not alter male communicative behavior.

**Figure 6.**
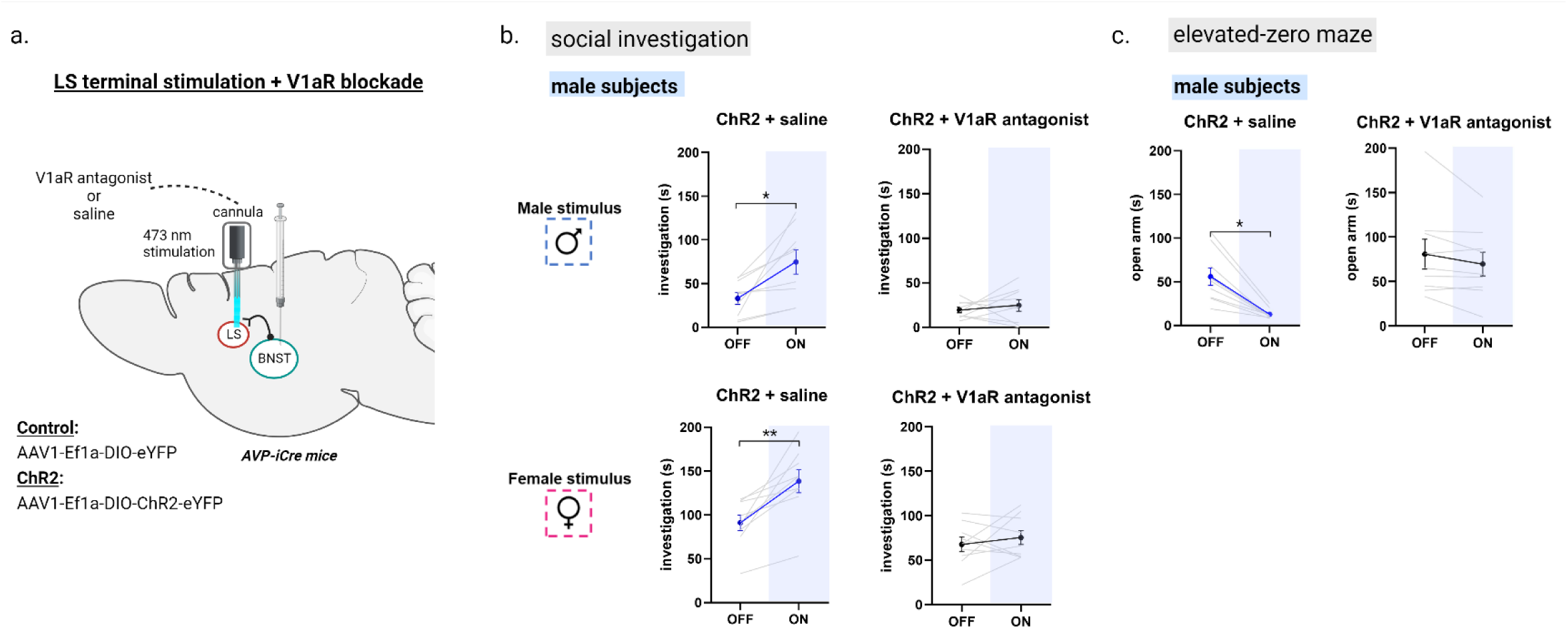
Antagonism of V1aR in the LS blocked optogenetic-mediated increases in male social investigation and anxiety-like behavior. (a) Bilateral BNST injection of ChR2 adeno-associated virus (ChR2) and fiber implantation within the lateral septum (intermediate zone). (b) Social investigation (in seconds) by male subjects during the three-chamber test. Two groups of ChR2+ injected males were tested with either a male or female stimulus (LIGHT OFF/ON), and received the same type of stimulus (i.e., novel male or female) with LIGHT OFF/ON + a highly-selective V1aR antagonist (subjects tested with a male stimulus: n=9; subjects tested with a female stimulus: n=9). All conditions were counterbalanced. Blue light stimulation (ON) of AVP-BNST-LS terminals in both ChR2 + saline male groups significantly increased time spent investigating male and female stimuli compared to investigation during light-OFF condition (Mixed model ANOVA, drug*light interaction, (F (1,16) = 11.76, p = 0.003; η^2^ = 0.42; *post hoc*: p = 0.01 (male stimuli), p = 0.006 (female stimuli)). In the same male subjects, antagonism of V1aR in the LS blocked optogenetic-mediated increases in male social investigation. (c) Time spent in the open arm of the elevated-zero maze (EZM). Blue light stimulation (ON) of AVP-BNST-LS terminals in ChR2 males significantly decreased time spent in the open arm of the EZM, and in the same males, antagonism of V1aR in the LS blocked optogenetic-mediated increases in male anxiety-like behavior. (Mixed model ANOVA, treatment*light interaction, (F (1,8) = 10.58, p = 0.012; η^2^ = 0.5). Bonferroni *post hoc* tests were used. *p<0.05, **p<0.01.

As was previously observed, BNST AVP → LS stimulation significantly decreased time spent in the open arm of the EZM, indicating this stimulation increased anxiety-like behavior (Figure 6c), but did not correlate with increased social investigation (Figure S6), indicating differences in mechanism. Surprisingly, given the role of the AVP-V1aR system in anxiety <u>(6)</u>, blocking V1aR in the LS during the light-off condition did not alter anxiety-like behavior (Figure 6c). However, blocking LS V1aR during BNST AVP → LS stimulation prevented stimulus-induced increases in anxiety-like behavior, suggesting that this effect is mediated by V1aR.

## Discussion

### Summary

Previous experiments had indicated that lesioning AVP cells of the BNST or inhibiting their AVP synthesis specifically affected social investigation in males but not in females. Unresolved was whether acute manipulation affects behavior in both sexes. Here we show that engaging in social investigation drives fos expression in BNST AVP cells in both sexes. Acute manipulation indicates that these cells appear to play a role in behavior in both sexes, but much more prominently so in males. In males, optogenetic inhibition of these cells reduced social investigation of other males but not females, while stimulation increased social investigation of males and females as well as communication toward females. In females, optogenetic inhibition did not affect social investigation, and stimulation only increased investigation of males. In a follow-up experiment we found that optogenetic stimulation of BNST AVP cell axon terminals in the lateral septum (LS) in males increased social investigation and anxiety-like behavior in the elevated-zero maze, but had no effect in females. These effects are likely mediated by V1aR since V1aR antagonism in the LS blocked the increases in male social investigation and anxiety-like behavior as well as optogenetically-induced modulation of LS cells. Our findings indicate that the sexually dimorphic AVP cells in the BNST contribute to sex-specific aspects of social approach and anxiety-like behavior through their connections with the LS, which are mediated by V1aR receptors.

### Limitations

Injecting cre-dependent viral vectors generated about 50% more cells expressing excitatory or inhibitory channel rhodopsins in males compared to females. This probably is directly related to males having more cells expressing AVP in BNST <u>(22)</u> and possibly to higher amounts of Cre expressed per cell. Lower levels of Cre expression per cell could in principle reduce the success rate of opsin expression per cell, which would lower the percentage of AVP cells in males and females that can be manipulated optogenetically. This does not appear to be the case as we find that the ratio of cells expressing opsins in males and females is similar to the normal ratio of AVP-expressing cells in BNST of these mice in an earlier study <u>(27)</u>. In addition, the level of opsin expression per cell likely does not appear to differ significantly in males and females; there are no differences in *ex vivo* (physiological responses to optogenetic stimulation/inhibition) and *in vivo* (Fos expression after ChR2-based excitation) cellular responses in males and females. In addition, in previous studies we were equally able to delete AVP cells in males and females with Cre-dependent activated caspase <u>(27)</u>, indicating that Cre-dependent viral vectors in our mice are equally effective in inducing gene expression in males and females. This suggests that our ability to excite or inhibit individual BNST AVP cells in males and in females does not differ and that, therefore, differences in the effect of such manipulation on behavior probably reflect differences in the number and behavioral impact of these cells.

Although optogenetic inhibition or excitation of BNST AVP cell bodies significantly affects social behavior, these studies do not identify whether these effects are caused by changes in AVP release from terminals. These cells are likely to produce several other transmitters, for example, galanin <u>(40)</u>, which has also been implicated in control of social behavior <u>(41)</u>. Previously, however, we have shown that blocking AVP synthesis in BNST cells reduce the same behaviors that are blocked by optogenetic inhibition in this study <u>(26)</u>, supporting the idea that changes in AVP release plays an important role in the changes observed in this study. An even stronger argument that changes in AVP release underlies these effects is that the effects of stimulating BNST → LS terminals could be blocked by AVP antagonists in this study.

We opted for testing the effects of local excitation of terminals in this set of projections, because previous research indicates an important role of AVP within the LS <u>(6, 9, 31)</u>. However, the behavioral effects of stimulating BNST AVP → LS terminals may be due to antidromic invasion of the soma that could increase activity at other BNST AVP projections. While we cannot completely exclude this possibility, the fact that blockade of V1a receptors in the LS prevented both ChR2-mediated excitation of the LS and male social investigation suggests that the LS is one of the major downstream targets for BNST AVP cells.

### Activity of BNST AVP cells

Our results indicate that social interactions increase fos expression in BNST AVP cells. Earlier studies have already shown that male mice involved in social interactions (aggression and copulatory behavior) had higher levels of Fos expression in BNST AVP cells in mice <u>(32)</u> as do males of several avian species <u>(25)</u>. Our study shows that even brief social contact with either sex can increase activity of BNST AVP cells in males as well as in females. However, there is a sex difference in the amount of Fos expression, with male cells showing higher percentage of Fos expression irrespective of stimulus. It is not clear whether this difference is related to intrinsic properties of cells or in the nature of the inputs these cells receive. Earlier we showed several sex differences in input, such that male BNST AVP cells receive more input from some regions, such as the medial amygdala, whereas females receive more input from other brain areas, such as the medial preoptic area <u>(30)</u>. These differences may contribute to the different roles these cells play in social behavior in males and females. At this point, however, it is unclear how or whether fos activation in these cells translates to changes in the output of these cells.

### Acute inhibition and excitation of BNST AVP cells

The effects of inhibiting BNST AVP cells on social investigation aligned very well with results of earlier studies where we specifically lesioned these cells or reduced their AVP synthesis <u>(26, 27)</u>. In males, all these manipulations reduced investigation of other males, but not of females, and, in females, they did not affect social investigation at all. Similar reductions of male-typical social proximity have been observed after AVP knockdown in the BNST of several bird species <u>(42–44)</u>. Excitation of these cells, however, affected social investigation in both sexes, although not necessarily in the same way. In males, it increased investigation of males as well as females, indicating that artificial stimulation of these cells can also drive male interest in females as well eliciting increased communicative behavior (urine marking) toward them. In females, stimulation effects were limited to an increase in social investigation of males; there was no increase in social investigation of other females. As females expressed less Fos in BNST AVP cells during social investigation, these cells may normally be less active during social behavior, perhaps due to sex differences in inputs <u>(30)</u>. Overall, these manipulations suggest a much more central role for BNST AVP cells in social investigative behavior and certain communicative behaviors in males than in females. In addition to the differences in inputs mentioned above, the output of these cells show important sex differences as well <u>(30)</u> and track the differences in density of vasopressin projections in areas, such as the LS, that receive steroid-sensitive AVP innervation (higher in males than in females) <u>(21, 45)</u>.

### Acute excitation of BNST AVP → LS on social investigation, anxiety-like behavior, and LS cell activity

Our results indicate that the BNST AVP → LS play a major role in BNST AVP control of social interest in males, paralleling AVP’s facilitatory action within the LS on social recognition <u>(5)</u>, pair-bonding and parental care <u>(46, 47)</u>, aggression <u>(48–50)</u>, social interactions <u>(51)</u>, and social communication <u>(52)</u>. Indeed, we observed a strong overlap in the effects of excitation of BNST AVP cell bodies and BNST AVP projection to the LS on male social investigation and urine marking, indicating that the LS is the major output structure regulating BNST AVP action on these behaviors. The stimulatory effects on investigation are probably mediated by AVP acting on V1aR-expressing cells in the LS, as these effects could be blocked by V1aR antagonists. Stimulatory effects on urine marking, however, could not be blocked by V1aR antagonists, suggest that AVP may exert its effects on urine marking via oxytocin receptors or V1b receptors, both of which are present in the LS <u>(48, 53)</u> and both of which can mediate central actions of AVP <u>(48, 54)</u>. Not all effects of BNST AVP projections on social behavior appear to be mediated by the LS. For example, in females, excitation of cell bodies stimulated social investigation, but excitation of BNST AVP → LS did not. It may be that BNST AVP projections to other areas other than the LS may play a more important role in the control of social behavior in females.

The fact that inhibition of AVP BNST cells affected social investigation in males only, whereas excitation affected it in both sexes, suggest that AVP BNST cells may chronically maintain a behavioral state more conducive for social investigation in males but not in females, perhaps by controlling anxiety or arousal states. In fact, central AVP has been consistently implicated in the generation of anxiety states <u>(31, 55)</u>, and AVP in the LS controls anxious states by acting on V1a receptors <u>(56–59)</u> and more prominently in males than in females <u>(35, 60)</u>. We were, therefore, surprised that we did not observe changes in anxiety-like behavior during BNST AVP cell inhibition or stimulation. However, when we stimulated BNST → LS AVP terminals, we did find an increase in anxiety-like behavior within the EZM that could be blocked with a V1aR antagonist, but only in males, consistent with prior research. If there is a link between the effects of manipulating BNST → LS AVP projections on anxiety and social behavior, this link is tenuous as we did not find a correlation between anxiety and social investigation in individual animals.

While it is clear that the BNST AVP system regulates male-typical social investigation, especially toward other males, the exact motivational state induced by this system’s activation is not clear. One possibility is that this system regulates anxiety-like states during stressful social encounters, as may occur when two males interact, which then influence the intensity and shape of social behaviors. It is also not clear what is gained by modulating the level of investigation. One possibility is that increasing the time spent investigating increases the quality of the evaluation of potential competitors during acute interactions <u>(61)</u> or allows for efficient formation of social memories, which appears to require this circuit <u>(62, 63)</u>.

Our electrophysiology backs the notion that BNST → LS AVP projections act on LS target cells via V1aR. Stimulation of these projections initially increases the firing frequency of these cells and then depresses them, changes that were not seen when the V1aR antagonist was present. These observations align with the bimodal effects of exogenous AVP on LS neurons, which causes an initial excitation in a subset of neurons, followed by a more generalized depression of most LS neurons <u>(64, 65)</u>. Identifying the inputs and targets of these AVP-responsive neurons may provide insight into how BNST → LS AVP projections control social behavior and anxiety states.

### Conclusions

Overall, our results suggest a sexually differentiated role for BNST AVP neurons with these neurons having a stronger impact on social investigative behaviors and anxiety-like behaviors in males than in females. A few other sexually differentiated neurochemical systems have been shown to play different roles in social behavior. For example, female-biased tyrosine hydroxylase neurons in the anteroventral periventricular nucleus (AVPV) drive maternal behavior in females and inhibit aggressive behavior in males <u>(66)</u>; male-biased BNST aromatase cells control social behavior in male, but not female mice <u>(67)</u> and female-biased cholecystokinin receptor-expressing cells in the VMHvl are critical for female, but not male sexual behavior <u>(68)</u>. In many of these cases, sexually differentiated systems drive behaviors in one, but not the other sex, even if these behaviors do not necessarily differ, such as anxiety-like behaviors in the present and other studies <u>(69)</u>. In other words, the neurochemistry underlying these behaviors differs by sex. Conversely, sex differences in neurochemical systems do not necessarily translate in sex differences in behavior <u>(70)</u>. Such sex differences have been called latent sex differences (e.g., <u>(71)</u>), and probably contribute to sex differences in vulnerabilities for behavioral disorders. The clear sex differences in BNST AVP cells and their connections offer an excellent opportunity to unravel the functional consequences of sex differences in neuronal structure.

## Materials and Methods

### Animals and Husbandry

All mice were maintained at 22°C on a 12/12 hr reversed light/dark cycle with food and water available *ad libitum*, housed in individually ventilated cages (Animal Care Systems, Centennial, CO, USA), and provided with corncob bedding, a nestlet square, and a housing tube. All animal procedures were performed in accordance with Georgia State University Institutional Animal Care and Use Committee regulations and the National Institutes of Health Guide for the Care and Use of Laboratory Animals.

### Subject Animals

Founding AVP-iCre mice were obtained from Dr. Michihiro Mieda (Kanazawa University, Japan). These mice were generated using a bacterial artificial chromosome that expressed codon-improved Cre recombinase under the transcriptional control of the AVP promoter (AVP-iCre mice). In these animals, iCre expression is found in the bed nucleus of the stria terminalis (BNST) and the medial amygdala (MeA), as well as in hypothalamic areas <u>(72)</u>. Subjects were derived by crossing heterozygous iCre mutants to wildtype C57Bl/6J mice and genotyped (ear punch) by polymerase chain reaction (PCR) at 21–24 days of age (Transnetyx). A total of 170 adult iCre+ mice (2-4 months old) were used for *in vivo* behavioral experiments (91 males, 79 females): Fos mapping: n = 15 males, n = 15 females; BNST AVP cell stimulation: n = 18 males, n = 22 females; BNST AVP cell inhibition: n = 20 males, n = 21 females; BNST AVP → LS stimulation: n = 20 males, n = 21 females; BNST AVP → LS stimulation with V1aR antagonist: n = 18 males. AVP-iCre+ mice (BNST AVP cell stimulation: n = 8 cells (males), n = 8 cells (females); BNST AVP cell inhibition: n = 16 cells (males and females); BNST AVP → LS stimulation: n = 7 cells (males); BNST AVP → LS stimulation with V1aR antagonist: n = 5 cells (males)) were used for *ex vivo* electrophysiological recordings. All subject mice were socially experienced (see below) and singly housed for a minimum of one week prior to experimental use.

### Stimulus Animals

CD1 (ICR; Charles River Laboratories, Wilmington, MA, USA) mice were used as stimuli for behavioral testing and to provide male and female subjects with social experience because strain differences between subjects and stimulus mice increase social investigation. Mice were used at 9–16 weeks of age and were novel to the subject to which they were exposed. Female stimulus mice were grouped-housed, ovariectomized, and hormonally primed with subcutaneous injection of estradiol benzoate (E; 5 μg/0.1 mL sesame oil) followed by subcutaneous injection of progesterone (P; 250 μg/0.1 mL sesame oil) 44-48 hr later and approximately 4 hrs prior to use. Female stimulus mice were given two sexual experiences before use in social experience and behavioral testing of subjects.

Two groups of stimulus males were used for social experience and behavioral testing. Males that were used as subordinate mice to provide aggressive experience to subjects were grouped-housed, gonadectomized (GDX), and subjected to two aggressive encounters with a dominant male (see below). Mice in the second group, which provided sexual experience to female subjects and were used as stimulus animals in the three-chamber social task, were singly housed, gonadectomized, implanted with testosterone (GDX + T), and then given two sexual experiences before use in social experience/behavioral testing.

### Viral Vectors

#### Fos mapping

AVP driven-, Cre-expressing BNST neurons were induced to express eYFP (AAV-EF1a-DIO-eYFP; serotype 5; Dr. Karl Deisseroth; Addgene_27056) for visualization of BNST AVP neurons.

#### Cell inhibition

AVP driven-, Cre-expressing BNST neurons were induced to express the blue-light activated inhibitory opsin, *guillardia theta* anion-conducting channelrhodopsin (stGtACR; hSyn1-SIO-stGtACR2-FusionRed; serotype 1; RRID:Addgene_105677) or eYFP alone as a control (AAV-EF1a-DIO-eYFP; serotype 5; Dr. Karl Deisseroth; RRID:Addgene_27056).

#### Cell and terminal excitation

AVP driven-, Cre-expressing BNST neurons were induced to express the blue-light activated excitatory opsin, channelrhodopsin-2 (ChR2) fused to eYFP (*in vivo*) or mCherry (*ex vivo*): AAV-EF1a-double floxed-hChR2; serotype 5; eYFP: RRID:Addgene_20298, mCherry: RRID:Addgene_20297) or eYFP alone as a control (AAV-EF1a-DIO-eYFP; serotype 5; Dr. Karl Deisseroth; RRID:Addgene_27056).

### Surgical Procedures

All surgeries were conducted using 1.5–3% isoflurane gas anesthesia in 80% oxygen and 20% nitrous oxide; 3 mg/kg of carprofen (subcutaneous) was given before surgery to reduce pain.

### Stereotaxic Surgery

Mice were positioned in a stereotaxic frame (David Kopf Instruments, Tujunga, CA, USA) with ear and incisor bars holding bregma and lambda level. After a midline scalp incision, a hand-operated drill was used to make holes in the skull, exposing the dura. For all subjects, 200 nL of viral vectors (see above) were delivered bilaterally to the BNST (coordinates: DV: -4.3, AP: +0.13, ML: ±0.8) at a rate of 100 nL/min using a 5-μL Hamilton syringe with a 30-gauge beveled needle mounted on a stereotaxic injector. Following virus delivery, the syringe was left in place for 5 min before slowly withdrawing it from the brain. For *in vivo* optogenetic stimulation, dual optic fiber cannulas (Doric Lenses, DFC_200/230-0.48_4mm_DF1.6_DFL) were chronically implanted immediately following viral injections using the same coordinates for cell manipulation experiments or above the lateral septum (coordinates: DV: -3.0, AP: +0.35, ML: ±0.35) for terminal stimulation experiments. For dual-delivery of V1aR antagonist and light stimulation in the lateral septum, Dual Optofluid cannulas (Doric Lenses, DFC_200/230-0.48_3.5mm_DF0.8_DFL) with interchangeable injectors were implanted. All implants were secured to the scalp with one bone screw, one layer of dental cement (Calk Dentsply) and two layers of Ortho-jet powder and liquid mix (Lang Dental Manufacturing Co Inc). To allow for optimal viral expression, mice were allowed to recover for at least 14 days (cell stimulation) or 6 weeks (terminal stimulation) prior to use.

### Gonadectomy and Hormone Treatment (stimulus animals)

Testes were cauterized and removed at the ductus deferens via a midline abdominal incision. Silastic capsules (1.5 cm active length; 1.02 mm inner diameter, 2.16 mm outer diameter; Dow Corning Corporation, Midland, MI, USA) were filled with crystalline testosterone (T; Sigma, St. Louis, MO, USA) and inserted subcutaneously between the scapulae after gonadectomy (GDX + T); this procedure leads to physiologic levels of T. To further reduce aggression in stimulus animals, some males were gonadectomized, but did not receive a T implant (GDX). The ovaries of stimulus female mice were removed by cauterization at the uterine horn and attendant blood vessels.

### Social Experience

As opposite-sex sexual experience and attaining competitive status (“social dominance”) promote male and female social and communicative behaviors <u>(73)</u>, subject mice received adult social experience. This experience consisted of two of the following sequences: an opposite-sex encounter (sexual experience) followed by a same-sex encounter (aggressive experience) the following day with at least one day between social experiences.

### Opposite-sex (sexual) Experience

Subjects were given two opportunities to interact with either a stimulus female (for male subjects) or a stimulus male (for female subjects). A sexually experienced stimulus mouse was placed in the subject’s home cage and removed the next day (overnight, first sexual experience) or after 90 min (second sexual experience). Subjects that did not engage in any sexual behavior (mounting, intromission, or ejaculation) during the second sexual experience were removed from further testing.

### Same-sex (aggressive) Experience

Male subjects were exposed to two interactions with subordinate males (GDX) treated with 40 μL of GDX + T stimulus male urine applied to their backs. Gonadectomy, group housing, and social defeat of subordinates reduce offensive aggression in mice, while GDX + T male urine provides subjects with a male urinary cue that elicits offensive aggression. Subordinate stimulus males were placed in the subject’s home cage and removed after the subject’s first offensive attack (biting) within a 10-min period. All subject males attacked the intruder male stimulus by the second encounter, and all subordinate stimulus males displayed submissive behavior, defined as defensive postures (e.g., on-back), fleeing, and non-social exploring. Female subjects were exposed to a female intruder for a 10-min period; however, this did not elicit any attacks from either animal.

### Experimental Procedures (In vivo)

All testing occurred during the dark phase under red light illumination, except for the elevated zero maze (EZM) which was conducted during the light phase. Two-six weeks after viral injection and implantation surgery (or viral injection alone), subjects were habituated to the testing room and apparatus by handling and placing mice (for 3-5 min) in the three-chamber apparatus (see below) each day for 3 days. On experimental days, subjects were adapted to the experimental room for 15 min before testing. All behavioral tests were scored by an experimenter blind to the viral and drug manipulation of the subject.

To assess cellular activity in response to social stimulation, a group of AVP-iCre+ animals were injected within the BNST with Cre-dependent eYFP virus (see above) and exposed for 10 min to either a caged male, caged female, or an empty cage (with another adjacent empty cage) within a three-chamber apparatus. Mice were removed and killed 70 min later; brain tissue was extracted, sectioned, and processed for Fos/eYFP immunohistochemistry, with Fos as a marker for neural activity.

Subjects underwent a total of four tests for social communication and social approach in the three-chamber apparatus (62 (L) x 40 (W) x 22 (H) cm) with at least four days off in between test days. Each subject received two test days with light stimulation and two test days without light stimulation with each stimulus type (male and female live conspecifics). The order of treatment (light-ON, light-OFF) and stimulus condition (male, female) was counterbalanced across subjects, except that subjects exposed to a stimulus type on the first test were then given that same stimulus type on the second test with the opposite treatment condition. Female subjects were tested irrespective of estrous cycle day. Prior research indicates minimal effects of the estrous cycle on social investigation and communication <u>(74)</u>. Finally, mice were tested on either an EZM to evaluate their anxiety-like behavior or, in a subset of animals (n=9), for novel object investigation (Lego or Hot Wheel car) within a three-chamber apparatus. One week after behavior testing, subjects were sacrificed and perfused and had their brain tissue extracted and processed for immunohistochemistry to verify viral targeting of BNST. In addition, subjects that received BNST AVP cell ChR2-mediated stimulation first underwent 10 min of light stimulation in their home cage before being sacrificed and perfused 70 min later so that the level of activation (Fos colocalization) of ChR2-infected AVP cells could be determined.

### Light Stimulation

A dual fiber-optic patch cord (Doric Lenses, DFP_200/230/900-0.48_1m_DF1.6-2FC) was coupled, using a zirconia sleeve (Doric Lenses, SLEEVE_ZR_2.5), to dual optic fiber cannulas chronically implanted in subjects. During testing, the dual-fiber optic patch cord was connected via a FC/PC connector to a rotary joint (Doric Lenses, FRJ_1x2i_FC-2FC_0.22) to minimize torque on the animal’s head. The rotary joint was mounted above the center of testing arenas using a gimbal holder (Doric Lenses, GH_FRJ) to further minimize torsional stress on subjects during behavior testing. A mono-fiber optic patch cord (Doric Lenses, MFP_200/240/LWMJ-0.22_1m_FC-FC) connected the rotary joint to a 473 nm blue diode pumped solid-state laser (Shanghai Laser & Optics Century Co, BL473T8-150FC) via FC/PC connectors. Blue-light laser pulses were generated via a controller (PlexBright 4-Channel Optogenetics Controller) and Plexon Radiant Software (2.2.0.19). During ChR2 experiments, 10 Hz pulses of blue light (20 ms pulse width, 473 nm, 5-6 mW light power) were delivered at 5 s on/off intervals over a ten-minute (three-chamber tests) or five-minute period (EZM tests). Mice in the stGtACR2 cell inhibition experiment received constant light (473 nm, 5-6 mW light power) for the duration of each five-minute test. Light intensity optic fiber was verified prior to each test using a light sensor (Thor Labs, S140C); light power ranged from 5-6 mW at the tip of the dual optic fiber implant.

### V1aR Antagonist Injections into LS

The highly-specific V1aR antagonist (d(CH2)5[Tyr(Me)2,Dab5] AVP; Bachem) was diluted in sterile saline and 0.1% acetic acid to a final injected dose of 2.5 μM and stored at −20°C until use. This antagonist is exceptionally selective for V1aR, eliciting no detectable anti-OT activity *in vitro* or *in vivo* <u>(75)</u>. 30-45 min before behavioral testing, subjects were briefly anesthetized (1.5 - 3% isoflurane gas) and a 33 gauge needle was inserted through implanted guide cannulae, extending a total length of 3.7 mm. Subjects were then injected with 300 nL of V1aR antagonist or sterile saline (vehicle) at 100 nL/min (10 μL Hamilton syringe; Harvard Apparatus PHD 22/2000 syringe pump) via the guide cannula. The injection needle was left in place for 1 min to allow the drug to diffuse away from the tip of the injection needle, followed by replacement with optical fibers extending 3.2 mm.

### Social Investigation and Communication

Ultrasonic vocalizations (USV), urine marking, and social investigation were recorded in an acrylic three-chamber apparatus <u>(76)</u>. Instead of a solid floor, the apparatus was placed on absorbent paper (Nalgene Versi-dry paper, Thermo Fisher Scientific) to accurately measure urine marking. During testing with stimulus animals, subjects had access to a stimulus animal in a triangular cage [19 cm (hypotenuse), 15 cm (triangle legs), 22 cm (height)] and an empty (clean) cage placed at opposite corners of the outermost chambers of the apparatus. The location of the stimulus and clean cage were counterbalanced across subjects.

Subjects were placed in the center of the middle chamber after being connected to the dual-fiber optic patch cord and allowed to acclimate to the apparatus for 1 min. Animals were then allowed to investigate the apparatus for 10 min during which time close investigation of clean and stimulus cages, distance traveled throughout the apparatus, time spent in the stimulus and clean cage chambers, as well as USV production and urine marking were measured. After each test, the apparatus and cages were thoroughly cleaned with 70% ethanol and allowed to dry before further testing.

### Social Investigation and USVs

Close investigation was defined as time spent sniffing within 2 cm of the social stimulus or clean cage; climbing on the cage was not scored as investigation. USVs were detected using a condenser microphone connected to an amplifier (UltraSoundGate CM16/CMPA, 10 – 200 kHz, frequency range) placed directly above the center chamber. USVs were sampled at 200 kHz (16-bit) with target frequency set to 70 kHz (UltraSoundGate 116Hb, Avisoft Bioacoustics). Recordings were then analyzed using a MATLAB (MathWorks, RRID:SCR_001622) plug-in that automates USV analysis (MUPET; <u>(77)</u>. Sonograms were generated by calculating the power spectrum on Hamming windowed data and then transformed into compact acoustic feature representations (Gammatone Filterbank). Each 200 ms window containing the maximum USV syllable duration was then clustered, via machine learning algorithms, into USV syllable types (repertoire units) based on time-frequency USV shape. Repertoire units that appeared as background noise were discarded.

### Urine Marking

Following testing, the substrate sheet was allowed to dry for 1 hr and then sprayed with ninhydrin fixative (LCNIN-16; Tritech Forensics Inc.) to visualize urine marks <u>(78)</u>. After 24 h, sheets were imaged (Sony DSC-S700 camera), binarized and analyzed using a computer-aided imaging software (ImageJ, RRID:SCR_003070). Urine marking was measured both as the total area (cm^2^) and the number of visualized ninhydrin urine marks in the entire arena. During attachment of the patch cord to implants, subjects were restrained briefly which often elicited urination. Consequently, during cord attachment, subjects were placed over a separate sheet of absorbent paper so that urine pools produced during this procedure were counted separately from urine marks.

### Elevated Zero Maze (EZM) Test

The EZM consisted of a 5.5 cm wide circular platform (internal diameter 35 cm) raised 50 cm above the ground with two equally-spaced enclosed compartments and two open compartments. Subjects were connected to the patch cord and placed at the start of a closed compartment and allowed to explore the maze for 1-min prior to initiation of testing. During the 5-min testing period, subjects received either light stimulation (light-on) or no light stimulation (light-off) on separate tests, with the order of treatment (light-on, light-off) counterbalanced across subjects. Time spent in open and closed compartments of the EZM was manually scored from the recorded video.

### Real-Time Place Preference (RTPP) Test

To investigate whether BNST AVP cell stimulation/inhibition has inherent rewarding or aversive properties, subjects underwent a real-time place preference (RTPP) test. The test involved habituating the subjects to a clear plastic box divided into two equal-sized chambers (50 cm x 50 cm x 25 cm) and allowing them to explore freely for 1 min without any light stimulation. Subsequently, subjects were given a 10-min period to explore either a chamber paired with light stimulation or a chamber without light stimulation (randomized). Time spent in each chamber was recorded manually from which we generated a preference score to determine if light stimulation increased or decreased time spent in the light-associated chamber <u>(79)</u>.

### Histology and Immunohistochemistry

Animals were anesthetized with intraperitoneal injection of Beuthanasia-D (150 mg/kg) and transcardially perfused with PBS followed by 4% paraformaldehyde. Brains were extracted and post-fixed in 4% paraformaldehyde for approximately 12 hrs overnight. Brains were then transferred to a 30% sucrose solution and stored at 4°C before cryosectioning. Coronal sections were cut at a thickness of 30µm with a cryostat (Leica CM3050 S, Leica Biosystems) into 12-well plates filled with cryoprotectant and stored at -20°C before immunohistochemistry processing.

Sections were washed in 0.1M PBS (sequence of 5 washes for 5 min each) before being incubated with agitation overnight at room temperature with a primary antibody for eYFP/GFP (chicken anti-GFP; 1:5000; ab13970) and in 0.4% Triton-X in 0.1M PBS solution to detect AAV-infected cells. In addition, tissue from subjects in the ChR2 and Fos mapping experiments were also incubated with a primary antibody for cFos (rabbit anti-cFos, 1:1000, ab214672). The following day, sections were washed in 0.1M PBS again with the same sequence and incubated with agitation in the dark for 2 hrs in 0.4% Triton-X in 0.1M PBS with secondary antibodies: goat anti-chicken IgG (Alexa Fluor 488; 1:600; Invitrogen) and for Fos detection, goat anti-rabbit IgG (Alexa Fluor 594; 1:600; Invitrogen). Sections were washed in 0.1M PBS, mounted onto microscope slides (Fisherbrand Superfrost Plus), and cover-slipped using Prolong Gold (Invitrogen) for subsequent tissue analysis.

### Experimental Procedures (Ex Vivo)

#### Slice Preparation

Mice injected with either hSyn1-SIO-stGtACR2-FusionRed, AAV-EF1a-DIO-hChR2-mCherry, or AAV-EF1a-DIO-mCherry, were anesthetized with pentobarbital (50 mg/kg, IP) and then transcardially perfused with 20 ml of ice-cold sucrose artificial cerebrospinal fluid (aCSF) solution. This solution contained (in mM): 200 sucrose, 2.5 KCl, 1 MgSO_4_, 26 NaHCO_3_, 1.25 NaH_2_PO_4_, 20 D-glucose, 0.4 ascorbic acid, and 2.0 CaCl_2_, pH 7.2, 300-305 mOsmol/l. The mouse was then rapidly decapitated, and the brain was subsequently dissected, mounted in the chamber of a vibratome with superglue (Leica VT1200s, Leica Microsystems), and submerged into sucrose aCSF and bubbled constantly with 95% O_2_/5% CO_2_. Coronal sections containing LS or BNST were cut at a 300 μm thickness and placed in a holding chamber containing aCSF (in mM): 119 NaCl, 2.5 KCl, 1 MgSO_4_, 26 NaHCO_3_, 1.25 NaH_2_PO_4_, 20 D-Glucose, 0.4 ascorbic acid, 2 CaCl_2_, 2 Na^+^ Pyruvate and bubbled with 95% O_2_/5% CO_2_. Slices rested in a water bath at 25-32°C for 20 min to 1 hour before transfer to room temperature for recording.

#### Whole Cell Patch Clamp

For BNST AVP cell inhibition and excitation experiments, current clamp recordings targeted FusionRed/mCherry expressing cells were performed using an Axopatch 700B amplifier and a Digidata 1440A data acquisition system (Molecular Devices) on an Olympus upright microscope (BX51WIF) with a 40X water-immersion objective. For recordings in the LS, sections were placed on the stage of a Dragonfly 200 spinning disk confocal microscope system (Andor Technologies, USA) and neurons in close proximity to mCherry-labeled terminals projecting from the adjacent BNST were targeted for recording. Current clamp recordings were acquired using a MultiClamp 700A system and digitized with a Digidata 1440 (Molecular Devices, Sunnyvale, CA, USA).

Electrodes were pulled using a Flaming Brown horizontal puller (Sutter Instruments, Novato, CA, USA) from borosilicate capillaries (4-7 MΩ) and filled with internal solution (in mM): BNST AVP cell inhibition/excitation: K-gluconate 120, KCl 10, MgCl2 2, HEPES 10, CaCl2 0.1, EGTA 1, MgATP 3, and NaGTP 0.5 (pH 7.2 with KOH); LS terminal stimulation: 135 KMeSO_4_, 8 KCl, 10 HEPES, 2 Mg-ATP, 0.3 Na-GTP, 6 Phosphocreatine, as well as [mM] AlexaFluor 488 for visualization of neurons after recording (7.2 pH; 285-295 mol/kg H_2_O); the resistance of recording pipettes was 2-3 MW after being filled with internal solution. Once whole cell configuration was achieved, the cell was allowed to rest for 3 min to ensure stability of the patch and equilibration between internal solution and cytosol. For BNST AVP cell excitation, light stimulation of patched cells was driven by a blue-light laser (Ikecool Corporation) at 5Hz, 10 Hz, 20Hz or 40Hz lasting 5s or 90s. Spike fidelity, the accuracy with which neurons generated action potentials in response to light pulses, was also measured. For testing light-induced BNST AVP cell inhibition, current (10-50 pA) was injected to elicit action potentials (2-3 Hz) prior to, and during, blue-light inhibition. To measure the effects of ChR2-mediated stimulation of BNST → LS, current-clamped LS cells were held at a resting membrane potential where firing was present but steady (∼0.2-1 Hz). Stimulation of ChR2-expressing terminals in LS was then driven by an Andor mosaic blue-light laser using 5 s On/5 s Off pulse pattern at 10 Hz (20 ms pulse width, 80 ms interpulse interval) lasting 60 s. In all cases, cells that displayed a shift in series resistance that exceeded 20 MΩ or a 25% change from the start of the recording, or when the cell series resistance increased > 15%, were discarded from analysis.

Data was analyzed using ClampFit v10.7. Firing frequency was calculated from spiking data acquired in current clamp mode. Example traces were generated in Igor Pro 9 (WaveMetrics Inc., Portland, OR, USA). Statistics and graphs were generated with R or Prism (GraphPad, Boston, MA, USA).

#### Tissue Analysis

Bilateral images were taken at 10x magnification using a Zeiss Axio Imager.M2 microscope (Carl Zeiss Microimaging), which transferred fluorescent images (FITC contrast reflector) to image analysis software (Stereo Investigator, MicroBrightField, RRID:SCR_002526). Imaging domains (2 mm^2^) were placed with reference to anatomical landmarks (ventricles, fiber tracts; <u>(80)</u>). Subjects with off-target viral expression or implants were excluded from analyses. For Fos mapping and ChR2-mediated BNST AVP cell activation experiments, images taken under green fluorescence were overlaid with images taken under red fluorescence and cells were counted using ImageJ software (1.8.0_172). AVP cells expressing eYFP (green fluorescence), Fos (red fluorescence), or both (eYFP/Fos colocalization; yellow-orange) were counted bilaterally in the BNST across four sections/hemisphere, and averaged to determine the percent of cells co-expressing eYFP and Fos.

#### Statistical Analysis

All data were analyzed and graphed in R or Prism. All data met the assumptions of parametric statistical tests. Therefore, we analyzed *in vivo* data using mixed-model ANOVAs [between-subject factor: sex, virus (ChR2, eYFP); within-subject factors: light stimulation (OFF, ON), sex of stimulus (male, female)] followed by paired t-tests assessing treatment effects or one-way ANOVAs (Fos mapping experiment). All *post hoc* pairwise comparisons report Bonferroni-corrected p values and Cohen’s d for effect size when statistically significant. *Ex vivo* data was analyzed by using either one- or two-way repeated measures ANOVA. Results were considered statistically significant if p < 0.05.

## Data Availability

Behavior and histology data have been deposited in Figshare: https://figshare.com/s/b851f54722a244b4e8d4

## Acknowledgments and Funding Sources

This work was supported by the National Institutes of Health [R01MH121603; F31MH125659] and the Center for Behavioral Neuroscience at Georgia State University. Figures were created with BioRender.com.

