## Supplementary figures and images for "A vasopressin circuit that modulates sex-specific social interest and anxiety-like behavior in mice"

### Supplementary Figures

#### Slide 1
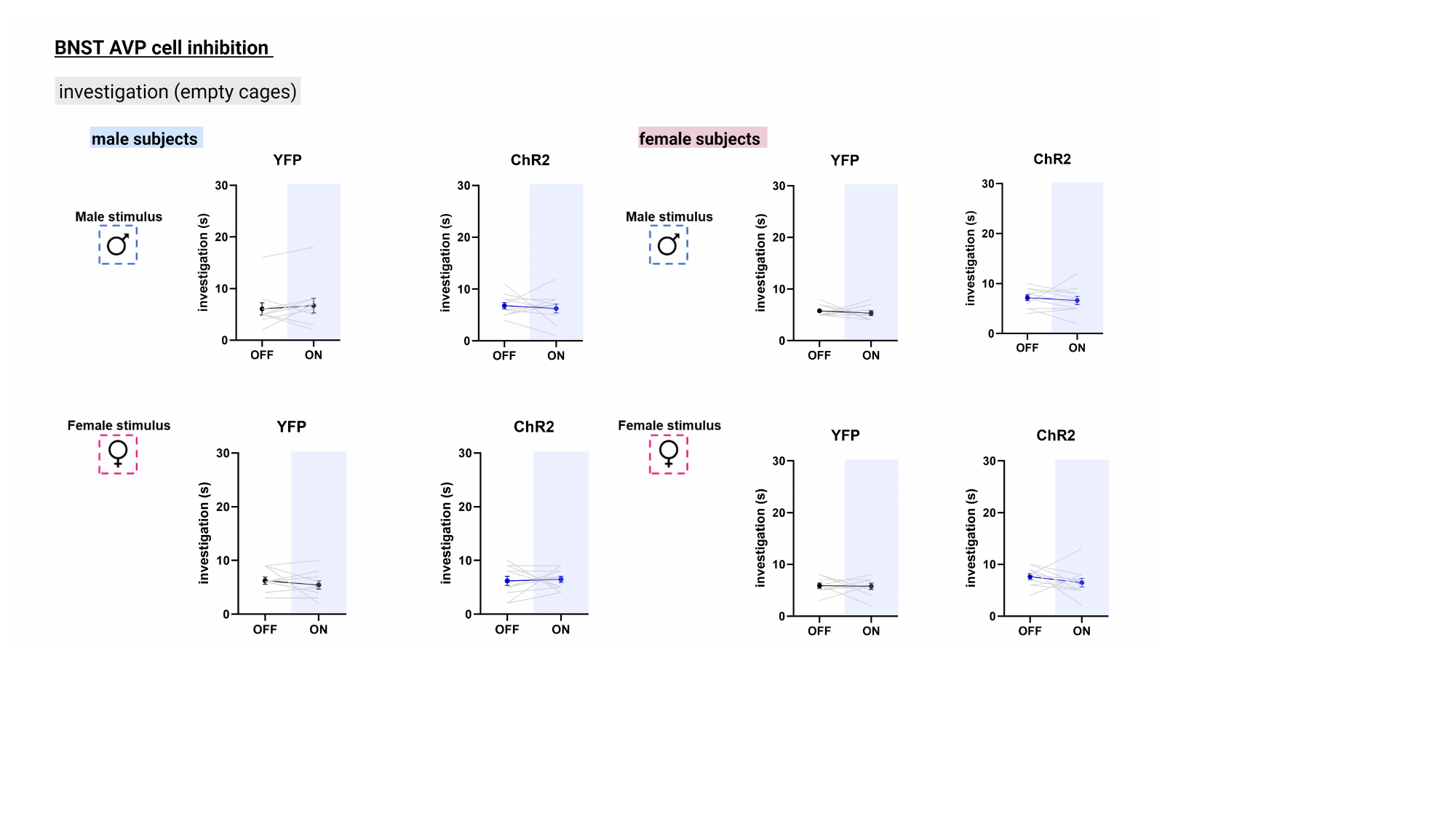

#### Slide 2
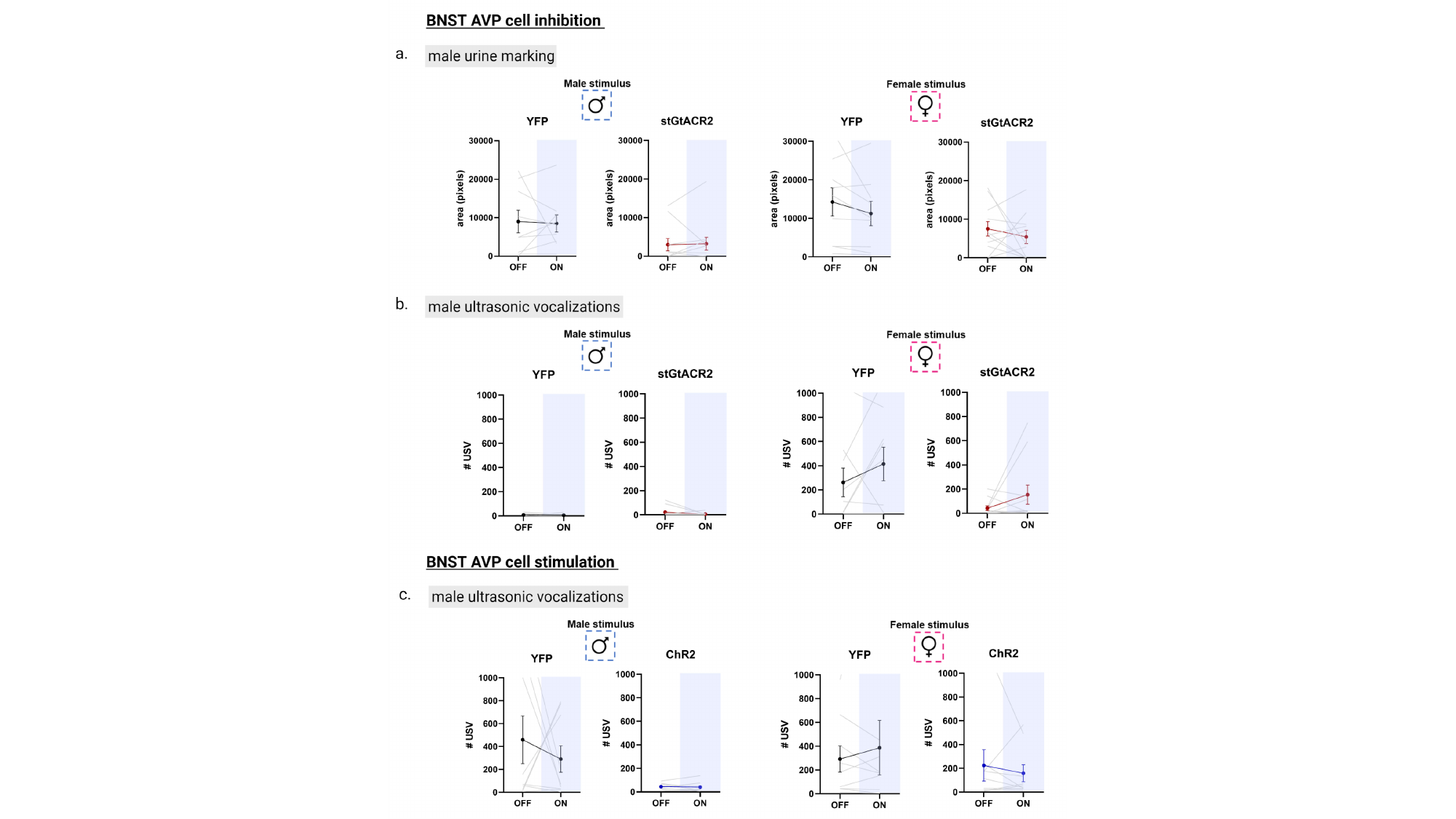

#### Slide 3
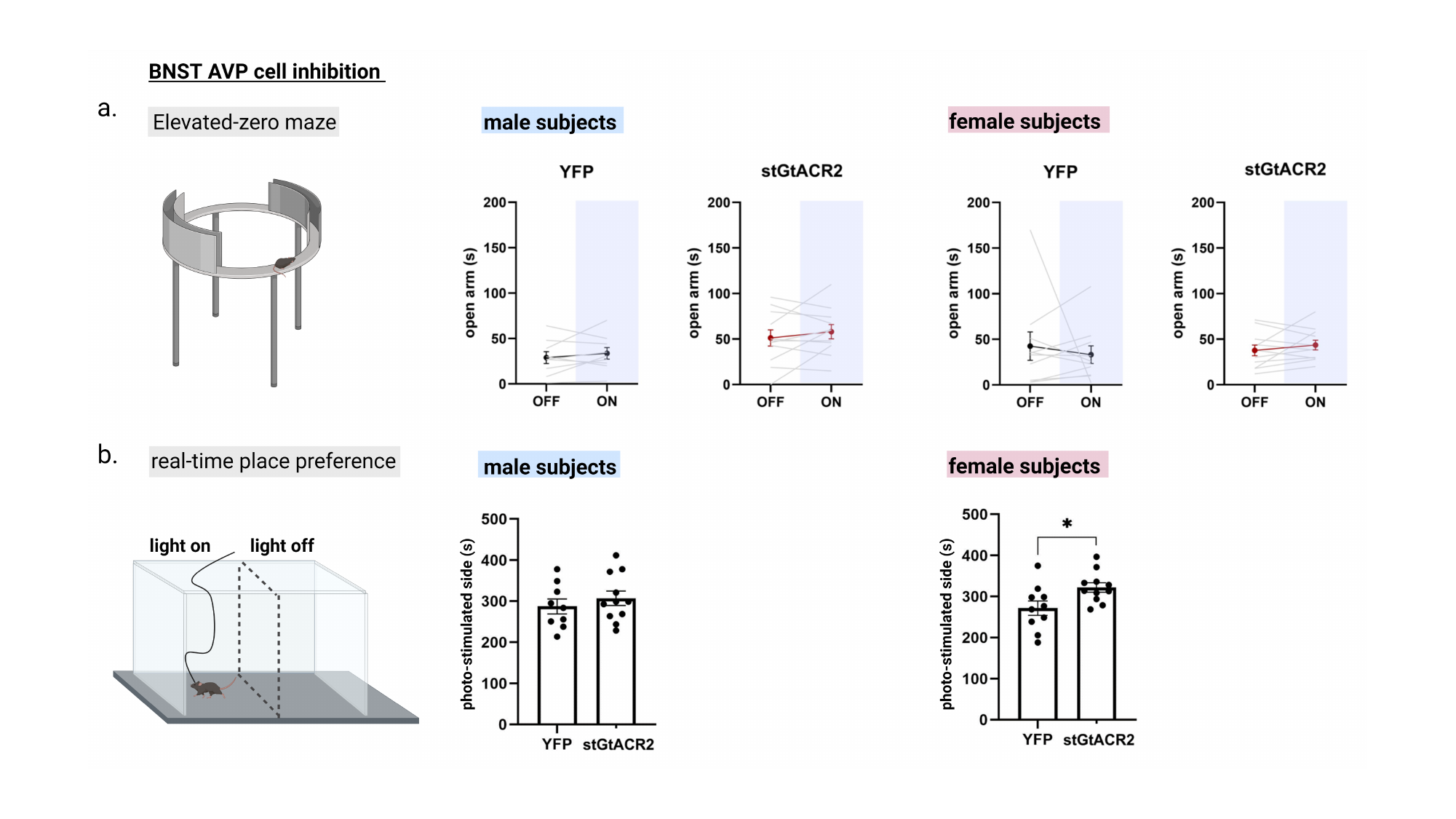

#### Slide 4
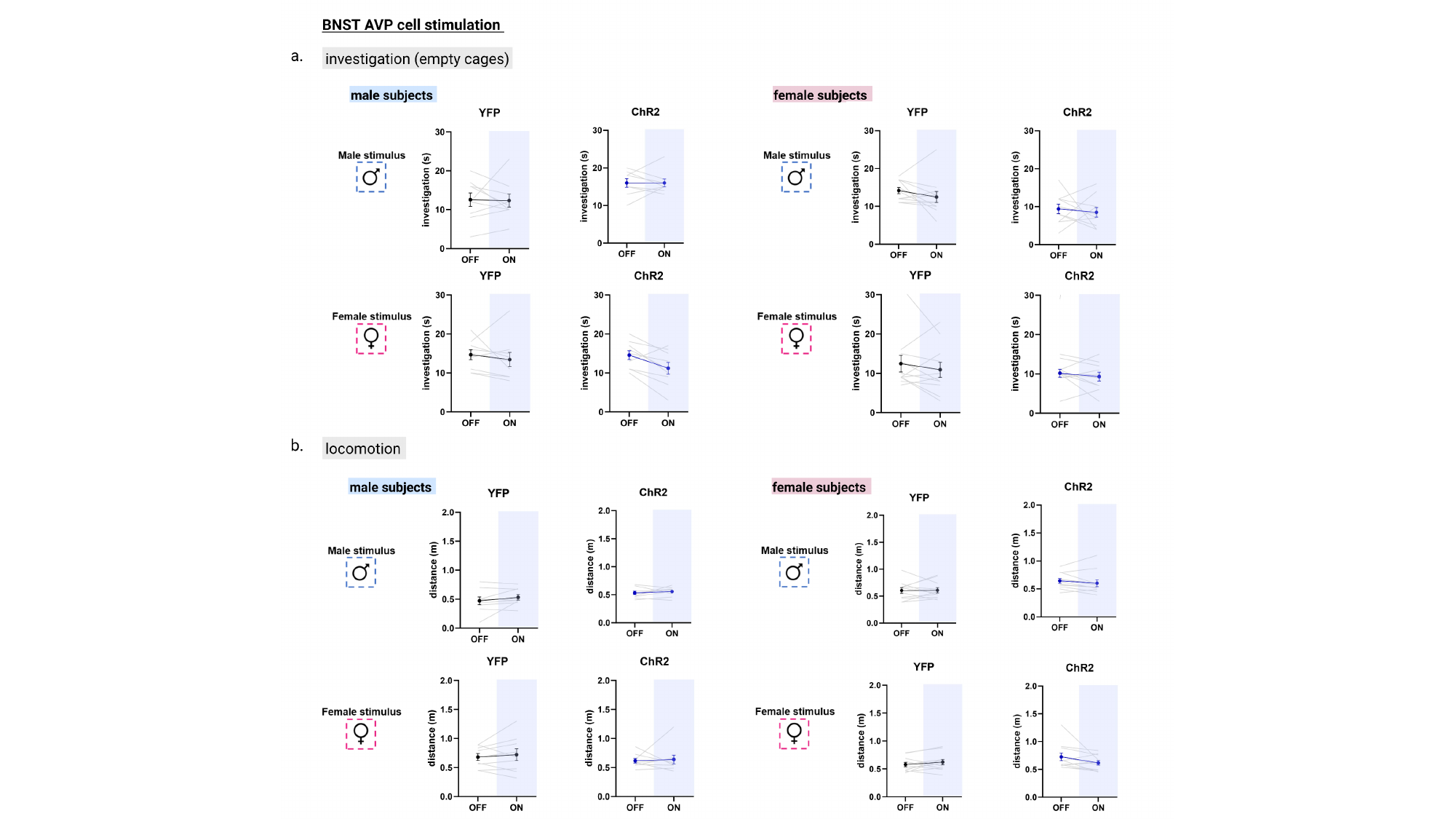

#### Slide 5
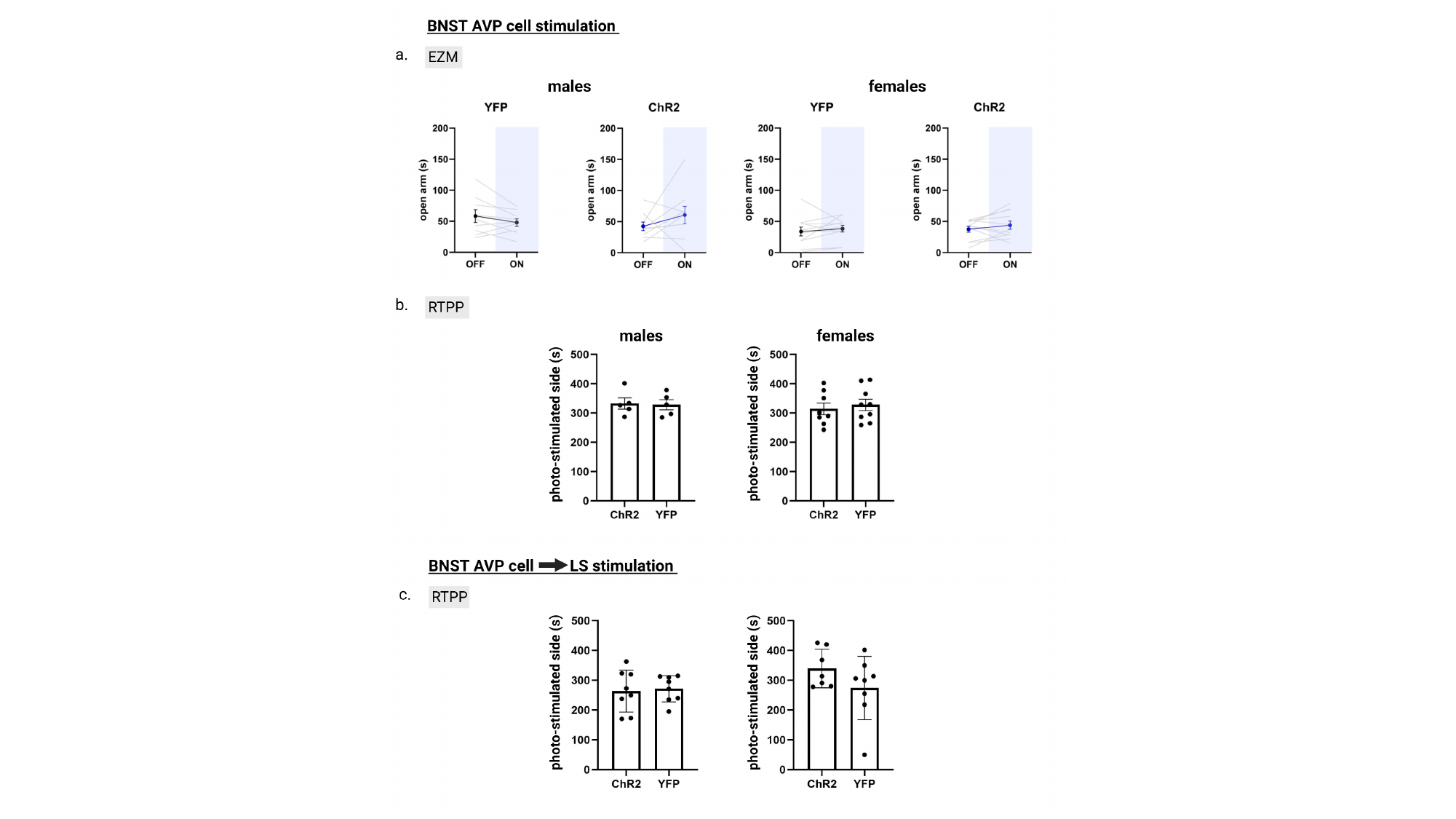

#### Slide 6
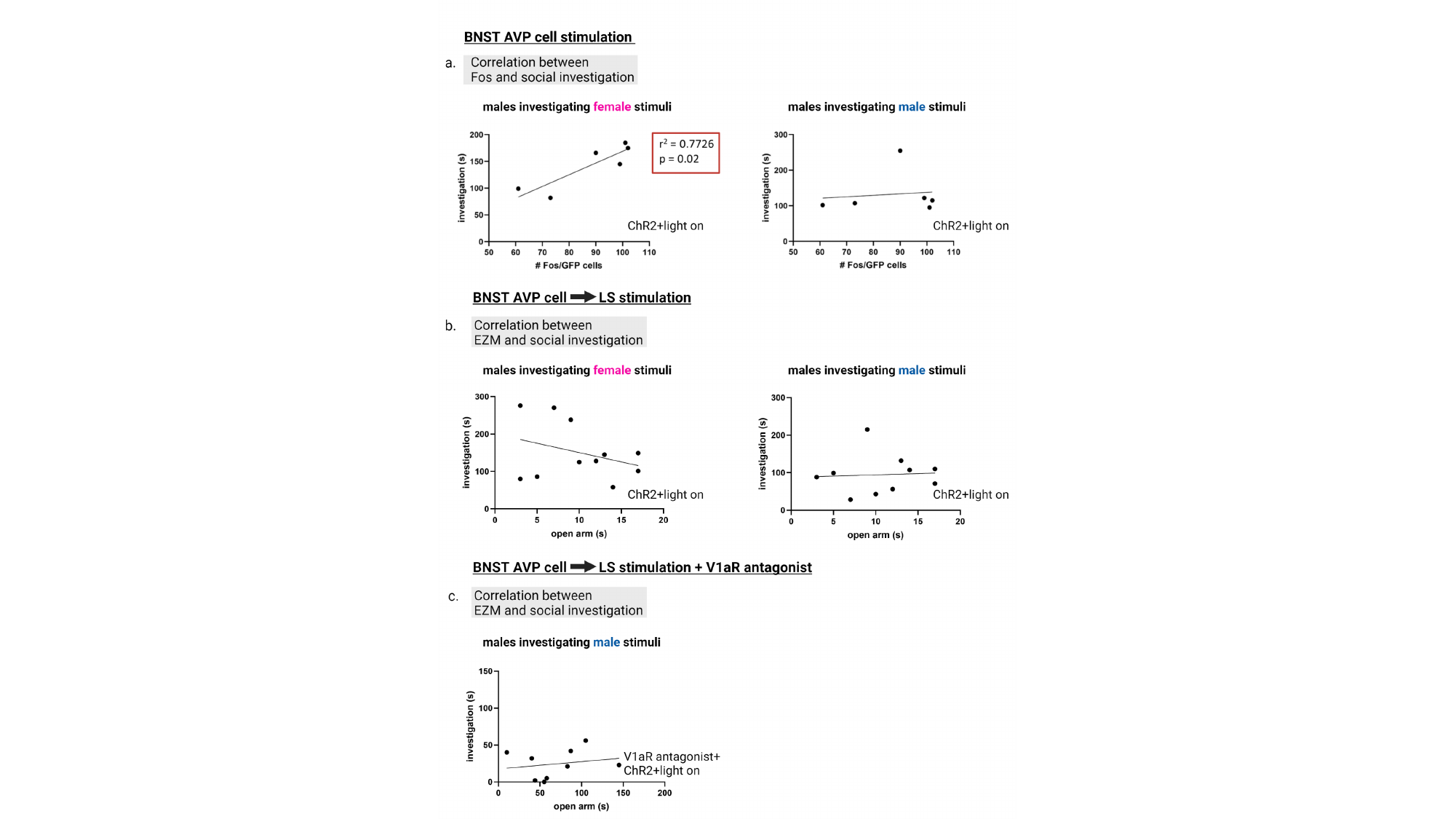

#### Slide 7
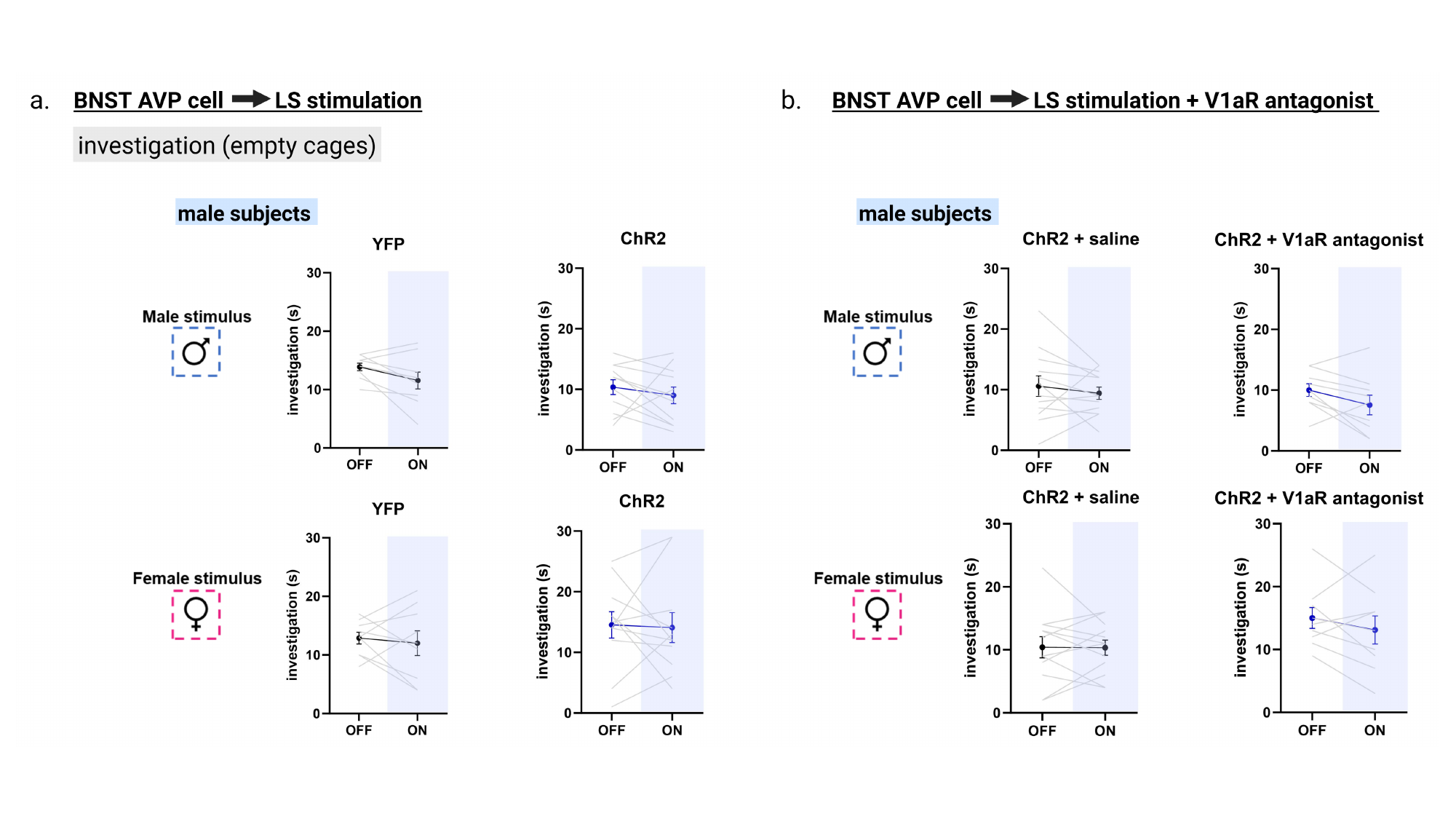

#### Slide 8
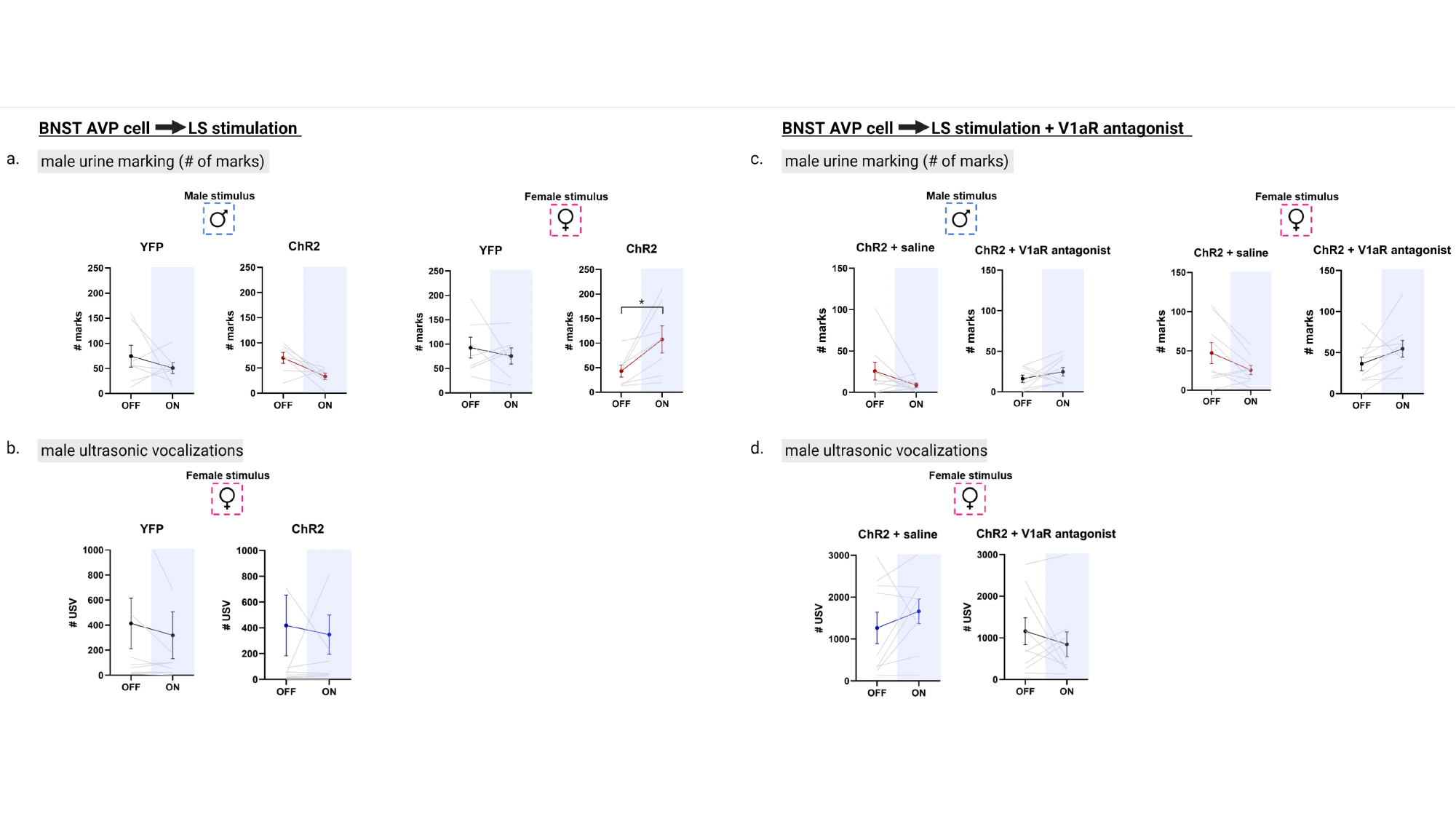

#### Slide 9
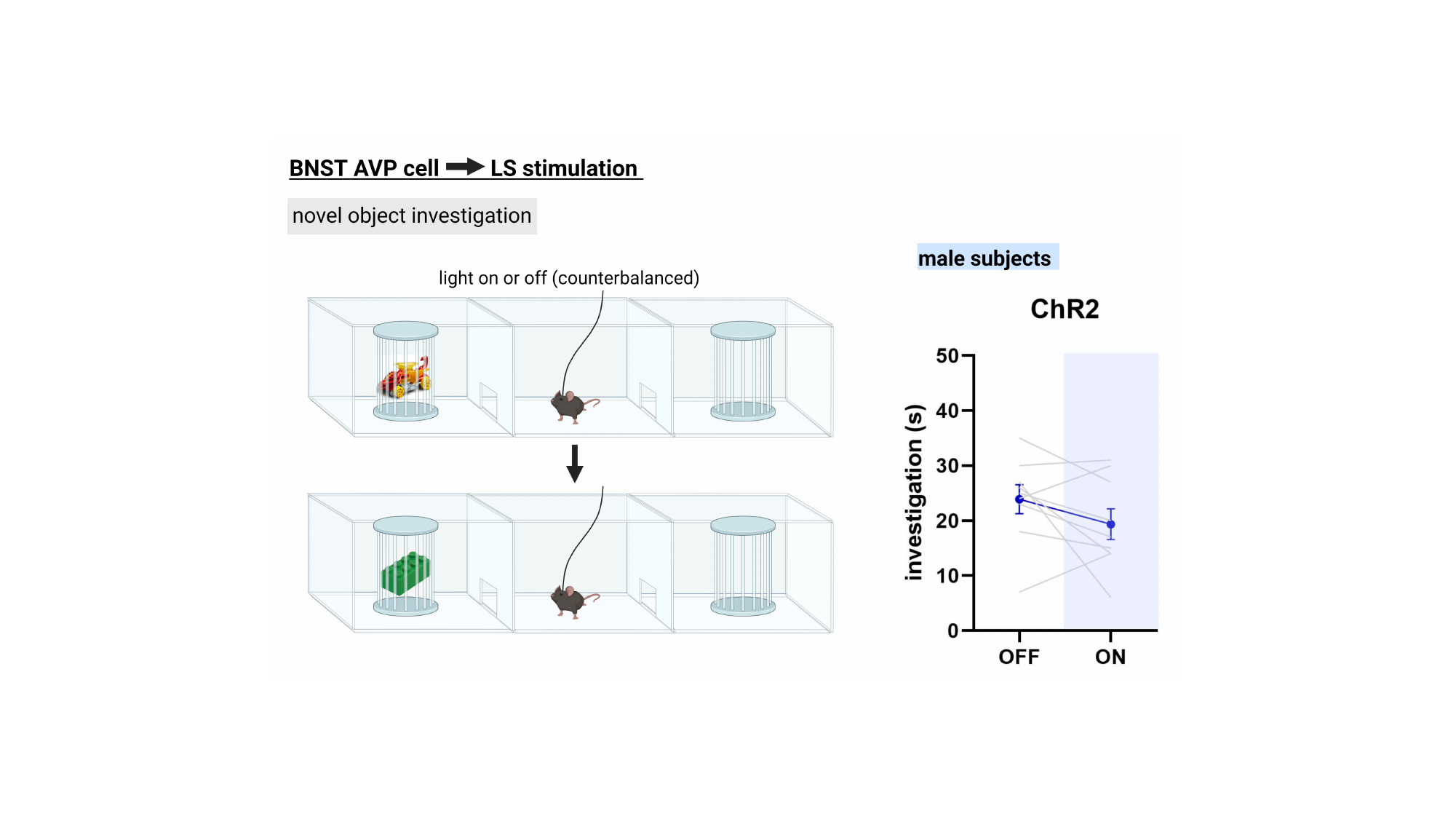

#### Slide 10
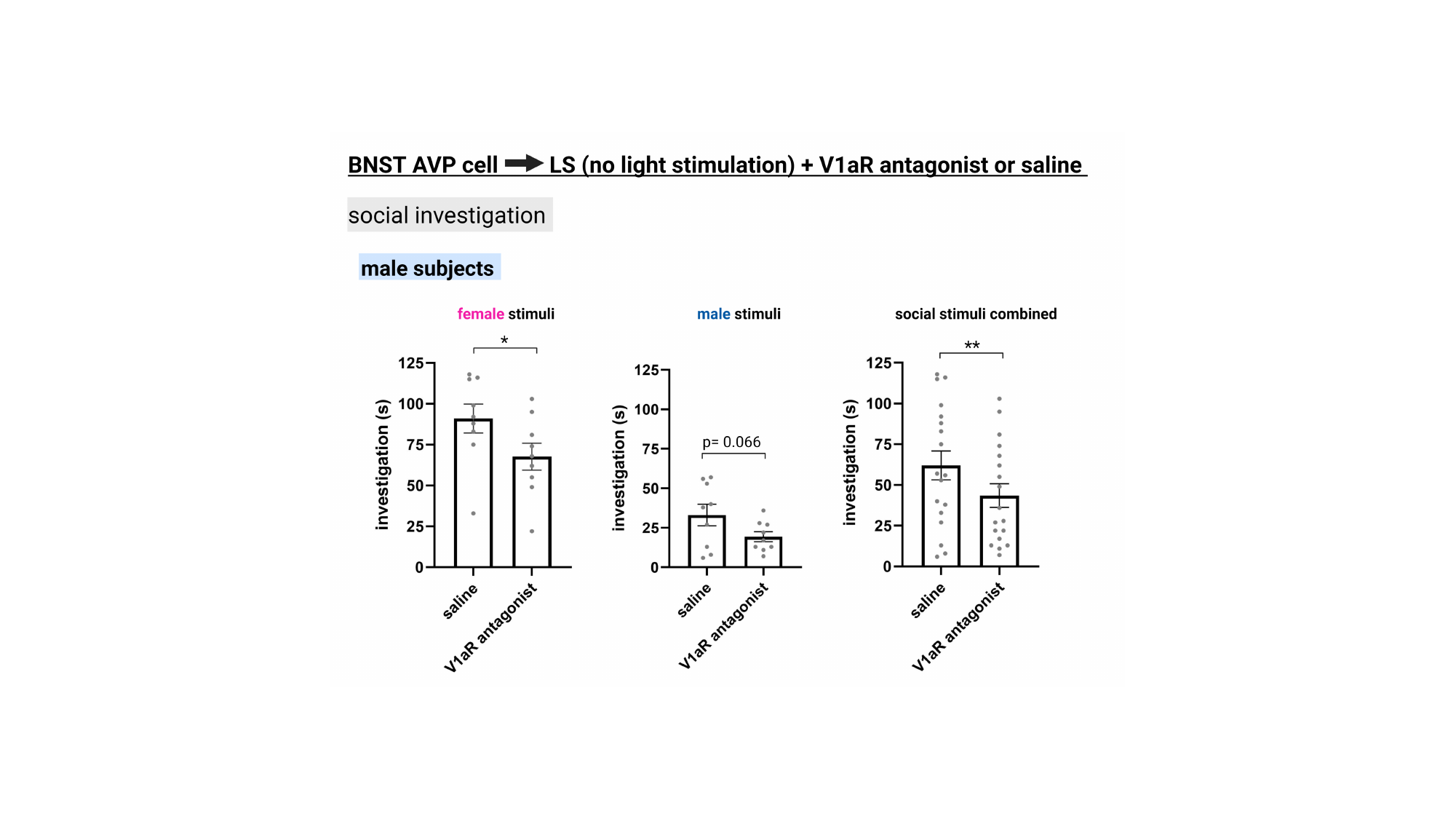
