## Supplementary Legends for "A vasopressin circuit that modulates sex-specific social interest and anxiety-like behavior in mice"

**Supplementary Figure 1** – Optogenetic activation of AVP-BNST cells did not alter investigation of empty cages during 3-chamber testing with a social stimulus or locomotion behavior. (a) Investigation of empty cages (in seconds) by male and female subjects during the three-chamber test (male subjects: YFP, n=9 and ChR2, n=9; female subjects: YFP, n=10 and ChR2, n=11) during light-OFF and light-ON conditions, counterbalanced. Light stimulation did not affect investigation times of YFP or ChR2 male and female subjects to empty cages in all 3-chamber testing. (b) Locomotion behavior measured by distance traveled (m) was not affected by light stimulation in all 3-chamber testing. Each point and horizontal line represent individual within-subject data. Overlapping data are represented as one point/line.

**Supplementary Figure 2** – BNST AVP cell manipulation did not affect male social communicative behaviors (urine marking, ultrasonic vocalizations. (a) In males, blue light inhibition (ON) of AVP-BNST cells in stGtACR2 and YFP controls did not affect urine marking or (b) ultrasonic vocalizations toward male or female stimuli. Females did not urine mark or produce ultrasonic vocalizations. (c) Optogenetic activation of BNST AVP cells did not affect ultrasonic vocalizations in males or females (females did not produce ultrasonic vocalizations). Each point and horizontal line represent individual within-subject data. Overlapping data are represented as one point/line.

**Supplementary Figure 3** – BNST AVP cell inhibition triggers real-time place preference in females, but not anxiety-like behavior (a) Time spent in the open arm of the elevated-zero maze (EZM). Blue light stimulation (ON) of stGtACR and YPF males and females did not alter time spent in the open arms of the EZM. (b) Real-time place preference. stGtACR females (BNST AVP cell inhibition) preferred to spend more time in the "light on" chamber compared to YFP control females (One-Way ANOVA, treatment*sex interaction, ($F(1,37) = 4.65$, $p = 0.038$, $\eta^2 = 0.11$; *post hoc*: $p = 0.024$). Light stimulation did not affect time spent in the "light on" chamber of ChR2 and YFP male subjects. Bonferroni *post hoc* tests were used. *$p<0.05$.

**Supplementary Figure 4 -** Optogenetic activation of AVP-BNST cells did not alter investigation of empty cages during 3-chamber testing with a social stimulus or locomotion behavior. (a) Investigation of empty cages (in seconds) by male and female subjects during the three-chamber test (male subjects: YFP, n=9 and ChR2, n=9; female subjects: YFP, n=10 and ChR2, n=11) during light-OFF and light-ON conditions, counterbalanced. Light stimulation did not affect investigation times of YFP or ChR2 male and female subjects to empty cages in all 3-chamber testing. (b) Locomotion behavior measured by distance traveled (m) was not affected by light stimulation in all 3-chamber testing. Each point and horizontal line represent individual within-subject data. Overlapping data are represented as one point/line.

**Supplementary Figure 5** – Elevated Plus Maze (EPM) and Real-time place preference (RTPP) was unaffected by BNST AVP cell stimulation and BNST AVP → LS terminal stimulation. (a) BNST AVP cell stimulation: time spent in the open arms of the elevated-zero maze (EZM). Blue light stimulation (ON) of BNST AVP cells in males and females did not affect time spent in the open arm of the EZM. (b) BNST AVP cell stimulation: Real-time place preference (RTPP). Blue light stimulation (ON) of BNST AVP cells in males and females did not affect time spent in the "light on" chamber of ChR2 and YFP subjects. (c) BNST AVP → LS terminal stimulation: Real-time place preference (RTPP). Blue light stimulation (ON) of AVP-BNST-LS terminals in males and females did not affect time spent in the "light on" chamber of ChR2 and YFP subjects.

**Supplementary Figure 6** – Correlations with social investigation. (a) The number of excited BNST AVP cells (Fos/YFP colocalized) in ChR2 males positively correlated with their time spent investigating female, but not male, stimuli ($r^2 = 0.7726$, $p=0.02$) (b) During AVP-BNST-LS terminal stimulation, time spent in the open arm of the EZM did not correlate with time spent investigating male or female stimuli. (c) During AVP-BNST-LS terminal stimulation + V1aR antagonism in the LS, time spent in the open arm of the EZM did not correlate with time spent investigating male stimuli.

**Supplementary Figure 7** – Optogenetic activation of LS terminals did not alter investigation of empty cages during 3-chamber testing with a social stimulus. (a) Investigation of empty cages (in seconds) by male subjects during the three-chamber test (LS male subjects: YFP, n=9 and ChR2, n=11) during light-OFF and light-ON conditions, counterbalanced. Light stimulation did not affect investigation times of YFP or ChR2 male subjects to empty cages in all 3-chamber testing. (b) Investigation (in seconds) by male subjects during the three-chamber test (subjects tested with a male stimulus: n=9; subjects tested with a female stimulus: n=9) during light-OFF and light-ON conditions, counterbalanced. Light stimulation did not affect investigation times of ChR2+saline or CHR2+V1aR antagonist male subjects to empty cages in all 3-chamber testing.

**Supplementary Figure 8** - AVP-BNST cell projections to the lateral septum (LS) and male social communicative behaviors (urine marking, ultrasonic vocalizations. (a) In males, optogenetic activation of AVP-BNST cell projections to the lateral septum (LS) increased the number of marks produced in the presence of a female in the 3-chamber apparatus (Mixed model ANOVA, treatment*light*stimulus interaction, ($F_{(1,17)} = 5.44$, $p = 0.038$; $\eta^2 = 0.4$; *post hoc*: $p = 0.03$) but did not affect (b) ultrasonic vocalizations toward female stimuli (males did not vocalize toward male stimuli). (b) Optogenetic activation of AVP-BNST cell projections to the lateral septum (LS) did not affect ultrasonic vocalizations in males or females (females did not produce ultrasonic vocalizations). (c) In males, optogenetic activation of AVP-BNST cell projections to the lateral septum (LS) with saline or V1aR antagonist administered did not affect the number of marks produced in the presence of male or female stimuli in the 3-chamber apparatus and did not affect (d) ultrasonic vocalizations toward female stimuli (males did not vocalize toward male stimuli). Females did not urine mark or produce ultrasonic vocalizations.

**Supplementary Figure 9** - Optogenetic activation of AVP-BNST cell projections to the lateral septum (LS) did not affect investigation of novel objects. Each subject received LIGHT OFF with a novel object (Hot Wheels car), then was tested again with LIGHT ON and another novel object (Lego) within the 3-chamber apparatus (males: n=9, novel objects, object side, and light condition counterbalanced).

**Supplementary Figure 10** – V1aR antagonism within the LS reduced male social investigation. Social Investigation (in seconds) by male subjects during the three-chamber test with no light stimulation (with a female stimulus: n=9; with a male stimulus: n=9; combined: n=18). Antagonism of V1aR in the LS decreased male investigation of female stimuli ($p=0.019$). Antagonism of V1aR in the LS caused a trend toward reduced male investigation of male stimuli ($p=0.066$). Antagonism of V1aR in the LS decreased overall social investigation compared to subjects that received control saline injections ($F(1,16) = 13.05$, $p = 0.002$, $\eta^2 = 0.45$). Bonferroni *post hoc* tests were used. *$p<0.05$, **$p<0.01$.
